# Human POT1 Prevents Severe Telomere Instability Induced by Homology Directed DNA Repair

**DOI:** 10.1101/2020.01.20.912642

**Authors:** Galina Glousker, Anna-Sophia Briod, Manfredo Quadroni, Joachim Lingner

## Abstract

The evolutionarily conserved POT1 protein binds single stranded G-rich telomeric DNA and has been implicated in contributing to telomeric DNA maintenance and the suppression of DNA damage checkpoint signaling. Here, we explore human POT1 function through genetics and proteomics discovering that the complete absence of POT1 leads to severe telomere maintenance defects that had not been anticipated from previous depletion studies. Conditional deletion of *POT1* in HEK293E cells gives rise to rapid telomere elongation and length heterogeneity, branched telomeric DNA structures, telomeric R-loops and telomere fragility. We determine the telomeric proteome upon POT1-loss implementing an improved telomeric chromatin isolation protocol. We identify a large set of proteins involved in nucleic acid metabolism that engage with telomeres upon POT1-loss. Inactivation of the homology directed repair machinery suppresses POT1-loss mediated telomeric DNA defects. Our results unravel as major function of human POT1 the suppression of telomere instability induced by homology directed repair.

## INTRODUCTION

Telomeres, the nucleoprotein structures at chromosome ends are distinct from chromosome internal breaks in remarkable ways. Telomeric proteins protect chromosome ends from degradation and suppress DNA damage signaling. In addition, they prevent DNA end-to-end fusions by non-homologous end joining and they suppress inter- and intra-chromosomal homologous recombination between telomere repeats, which can lead to rampant telomere elongation and loss events.

Many of the telomere functions are mediated by the shelterin proteins comprising TRF1, TRF2, RAP1, TIN2, TPP1 and POT1 which cover large parts of telomeric DNA (de Lange, 2018; Lazzerini-Denchi and Sfeir, 2016). POT1 is a highly conserved telomere binding protein which binds the telomeric single stranded G-rich DNA present as a 3’overhang at the end of the chromosomes (Baumann and Cech, 2001). Alternatively, when telomeres adopt a t-loop structure in which the single stranded 3’ overhang is tucked into double stranded part of the telomere (Doksani et al., 2013), POT1 is thought to bind to the single stranded displaced G-rich strand. Though POT1 binds directly to the single stranded telomeric DNA, it is recruited to telomeres through its physical interaction with the shelterin component TPP1 (Takai et al., 2011), which in turn interacts with TIN2 that is associated with the double strand telomere binding proteins TRF1 and TRF2 (de Lange, 2018). This type of molecular architecture is similar from mammals to fission yeast suggesting widespread conservation in evolution (Miyoshi et al., 2008).

Studies in a variety of eukaryotes unraveled important functions of POT1 in chromosome end protection and telomere length control. In fission yeast, Pot1 loss leads to rapid loss of telomeric DNA and chromosome circularization suggesting roles in protecting chromosome ends from nucleolytic degradation (Baumann and Cech, 2001; Pitt and Cooper, 2010). In mammals, such dramatic telomere loss events are not seen upon depletion of POT1. In contrast to the human genome, the mouse genome contains two POT1 paralogs, *Pot1a and Pot1b* and studies in this organism have contributed to the understanding of POT1 function (Hockemeyer et al., 2006; Wu et al., 2006). Deletion of conditional alleles in mouse embryonic fibroblasts (MEFs) revealed that POT1a represses ATR signaling which is triggered through RPA binding to single stranded telomeric DNA (Hockemeyer et al., 2006; Wu et al., 2006) whereas POT1b regulates the telomeric DNA overhang structure (Hockemeyer et al., 2006). *Pot1a* deletion in MEFs *per se* was also reported to increase homologous recombination (HR) in one study (Wu et al., 2006) but not the other (Hockemeyer et al., 2006). However, concomitant deletion of *Pot1a* and *Pot1b* induced higher incidences of telomere sister chromatid exchanges in *Ku70*^-/-^ cells suggesting contributions of mouse POT1a/b to the suppression of homology directed repair (HDR) in *Ku70*^-/-^ cells (Palm et al., 2009). The functions of POT1 in human cells had so far been mostly assessed by overexpression or RNAi-mediated depletion but not knockout studies. POT1-reduction in addition to the telomeric DNA damage response induced incorrect 5’ end processing in which telomeres lost the preferred ending with ATC-5’ to random positions within the telomeric repeat sequence (Hockemeyer et al., 2005).

Several studies also suggest direct or indirect roles of POT1 in semiconservative DNA replication and in telomere length control by telomerase. Overexpression of POT1 mutants which are prevalent in T cell lymphoma in human fibrosarcoma HT1080 cells increased telomere fragility which is indicative of DNA replication defects (Pinzaru et al., 2016). This effect was suspected to be due to defects of the POT1-mutants in POT1-assisted assembly of the CST complex (Pinzaru et al., 2016) which promotes lagging strand synthesis at telomeres. In addition, POT1 can regulate telomerase. Overexpression of a mutant form of human POT1 that lacks the DNA binding OB-fold induces rapid telomere elongation (Loayza and De Lange, 2003; Zhong et al., 2012) and POT1 also inhibits telomerase *in vitro* (Kelleher et al., 2005). On the other hand, in conjunction with TPP1, POT1 can stimulate telomerase processivity *in vitro* (Latrick and Cech, 2010).

In this paper, we assess human POT1 function by analyzing the consequence of POT1 loss upon conditional deletion in HEK293E cells. We demonstrate that the complete absence of POT1 triggers telomeric DNA branching, telomere elongation, accumulation of G-rich telomeric DNA, telomeric R-loops and telomere fragility. We elucidate changes in the telomeric proteome establishing an improved telomeric chromatin isolation protocol, identifying among others HDR factors that accumulate at telomeres in the absence of POT1. Significantly, the severe telomere maintenance defects but not the DNA damage response in the absence of POT1, are all suppressed upon inactivation of the HDR machinery. Thus, a major function of human POT1 is the repression of telomeric homologous recombination.

## RESULTS

### POT1 Loss Triggers Rapid Telomere Elongation and a DNA Damage Response Throughout the Cell Cycle

In order to study the functions of human POT1 for the maintenance of telomeric DNA structure and telomeric protein composition we genetically modified HEK293E cells. HEK293E cells were originally derived from a human embryonic kidney preparation but are presumably of adrenal origin (Lin et al., 2014). They contain the *E1A* adenovirus gene and *EBNA1* from Epstein Barr virus. We first tagged using CRIPSR/Cas9 gene editing the *TRF1* and *TRF2* genes at their N-termini with triple FLAG-tags to facilitate isolation of telomere chromatin as explained below. Cell clones were isolated and genotyped by PCR. Addition of these tags in chosen clone 293E cl75 did not interfere with telomere maintenance as assessed by telomere restriction fragment length (TRF) analysis nor did it induce a telomeric DNA damage response (Lin et al., manuscript in preparation). In a second step, we sought to develop a conditional allele of human *POT1* in 293E cl75, inserting by CRISPR/Cas9 gene editing LoxP sites in the introns flanking exon 6 (Figure 1A). We screened clonal isolates by PCR and sequencing for the appropriate genotype (Figures S1A and S1B). 293E cl75p100 carried two *POT1* alleles with the desired LoxP sites flanking exon 6 and a third allele with a frameshift mutation in exon 6 giving rise to a premature stop codon. Of note, HEK293E cells are triploid for chromosome 7, which harbors *POT1*. 293E cl75p100 maintain constant telomere length over several weeks of growth as assessed by TRF analysis (Figure S1C). We then transduced clone 293E cl75p100 with CreERT2 lentivirus expressing tamoxifen-dependent Cre recombinase to allow excision of the DNA between the LoxP sites. Cells were again cloned and screened for efficient editing of the conditional *POT1* alleles upon addition of 4-hydroxytamoxifen (4OHT) scoring for loss of POT1 protein expression by Western blot analysis (Figure S1D). Two clones (293E 75p100 Cre35 and 293E 75p100 Cre45) showed efficient loss of POT1 protein and induction of the DNA damage marker γ-H2AX as expected (Figure S1D). Phosphorylated forms of ATM and CHK1 also accumulated upon induced deletion of *POT1* (Figure 1C).

**Figure 1.**
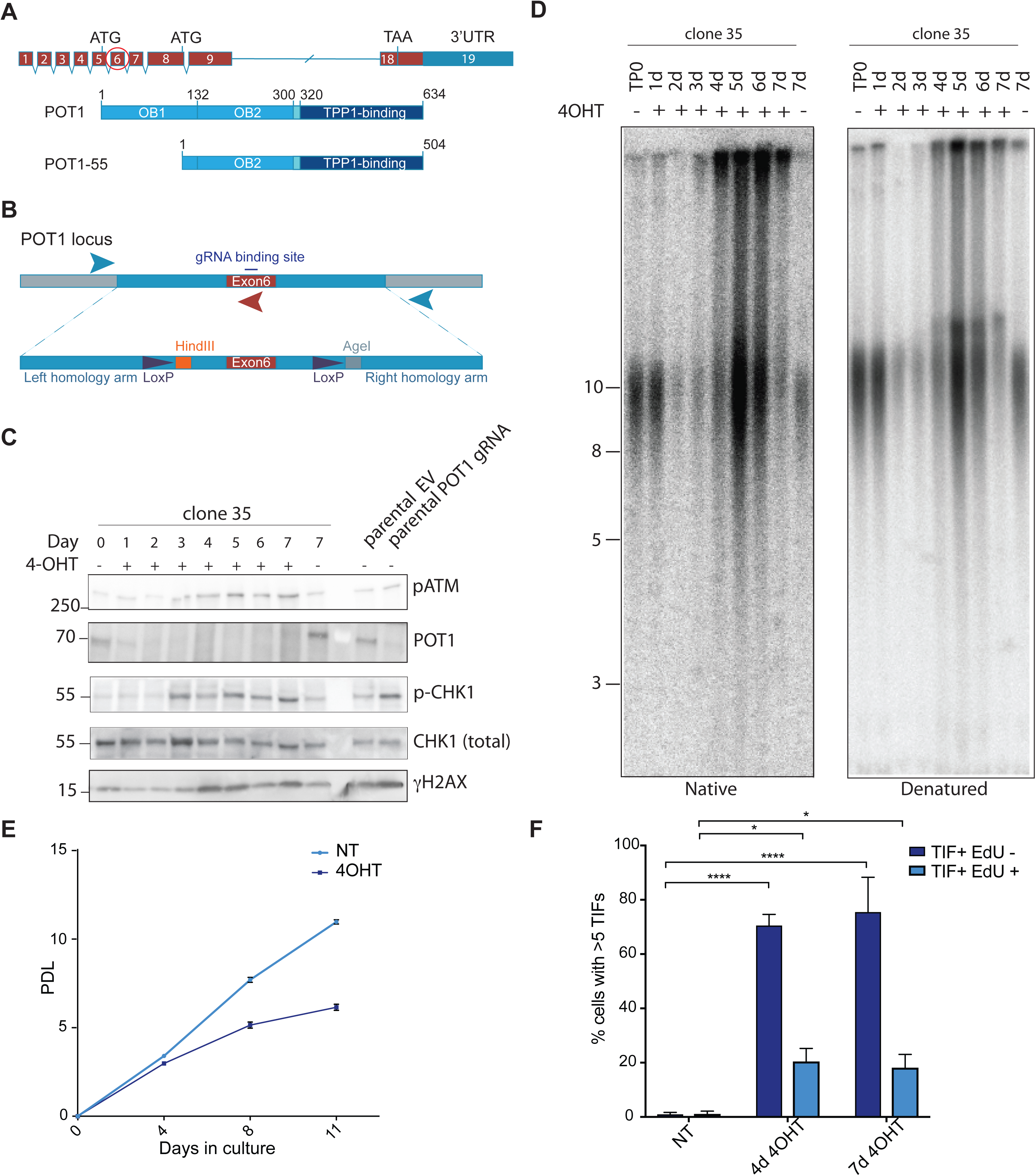
POT1 Loss Triggers Rapid Telomere Elongation and a DNA Damage Response Throughout the Cell Cycle. (A) Schematic drawing of the *POT1* gene structure and two derived polypeptides. Exon 6 that has been chosen for gene editing is encircled. (B) Schematic drawing of the repair template used for CRISPR/Cas9 mediated gene editing in HEK293E cells. Positions of primers for genotyping PCR are marked with arrows. (C) Western blot for POT1 and DDR markers upon *POT1* deletion induced with 0.5 μM 4-OHT. (D) Time course of telomere length changes upon POT1 removal in clone 35. d (days). (E) Growth curve for *POT1* WT and *POT1* knockout cells. Population doublings (PDL) are represented as mean ± SD. (F) Quantification for TIFs in EdU positive and EdU negative cells upon POT1 removal in clone 35. The bars show percentage of cells containing more than 5 TIFs + SD. Experiments were performed in triplicate. At least 100 cells were analyzed per condition per replicate. Significance was determined using two-way ANOVA. * p<0.05, **** p<0.0001

We assessed the effects of POT1 loss on telomeric DNA by TRF analysis of non-denatured DNA to detect single stranded G-rich telomeric DNA and of denatured DNA to measure telomere length distribution (Figures S1E and 1D). Very strikingly, loss of POT1 for more than 3-4 days caused dramatic telomere elongation, an increase of single stranded G-rich telomeric DNA and an accumulation of telomeric DNA molecules that could not enter the gel, suggestive of branched DNA structures. The viability and the proliferation of the cells upon POT1 loss were strongly affected after 4 days of culture in presence of 4OHT (Figure 1E). Analysis of cellular DNA content by fluorocytometry indicated that during the time course, the fraction of cells in G1 was reduced and a subset of cells with a sub-G1 DNA content accumulated indicative of cells undergoing cell death by apoptosis. The number of polyploid cells also increased indicating abnormalities in mitosis (Figure S2A). Immunofluorescence analysis of γ-H2AX foci showed that they colocalized with telomeric DNA in line with the crucial roles of POT1 in suppressing DNA damage specifically at chromosome ends (Figure S2B). We pulse-labeled cells with EdU to identify cells in S phase. The DNA damage foci were detected in both pulse-labeled EdU positive and EdU negative cells (Figure 1F), which lead us to conclude that POT1 suppresses DNA damage response throughout cell cycle.

### POT1 Loss Leads to Accumulation of Single-Stranded G-Rich DNA, Branched DNA Structures and Telomere Fragility

To test if the single stranded G-rich DNA at telomeres was terminal, we treated isolated genomic DNA with commercial 3’-5’ exonuclease I in vitro and repeated the native and denatured TRF analysis upon short-term gel electrophoresis which facilitates the quantification of the telomeric signals (Figure 2A). As expected, the G-strand DNA signal on native gels disappeared upon exonuclease I treatment in the uninduced cells containing wild type POT1 because the signal stems from the telomeric 3’ overhang only (Figure 2A). Upon deletion of *POT1* (lanes labeled with + 4OHT), the native signal was only partially reduced with exonuclease I, indicating that the telomeric DNA accumulated terminal as well as internal single stranded G-rich DNA in the absence of POT1.

**Figure 2.**
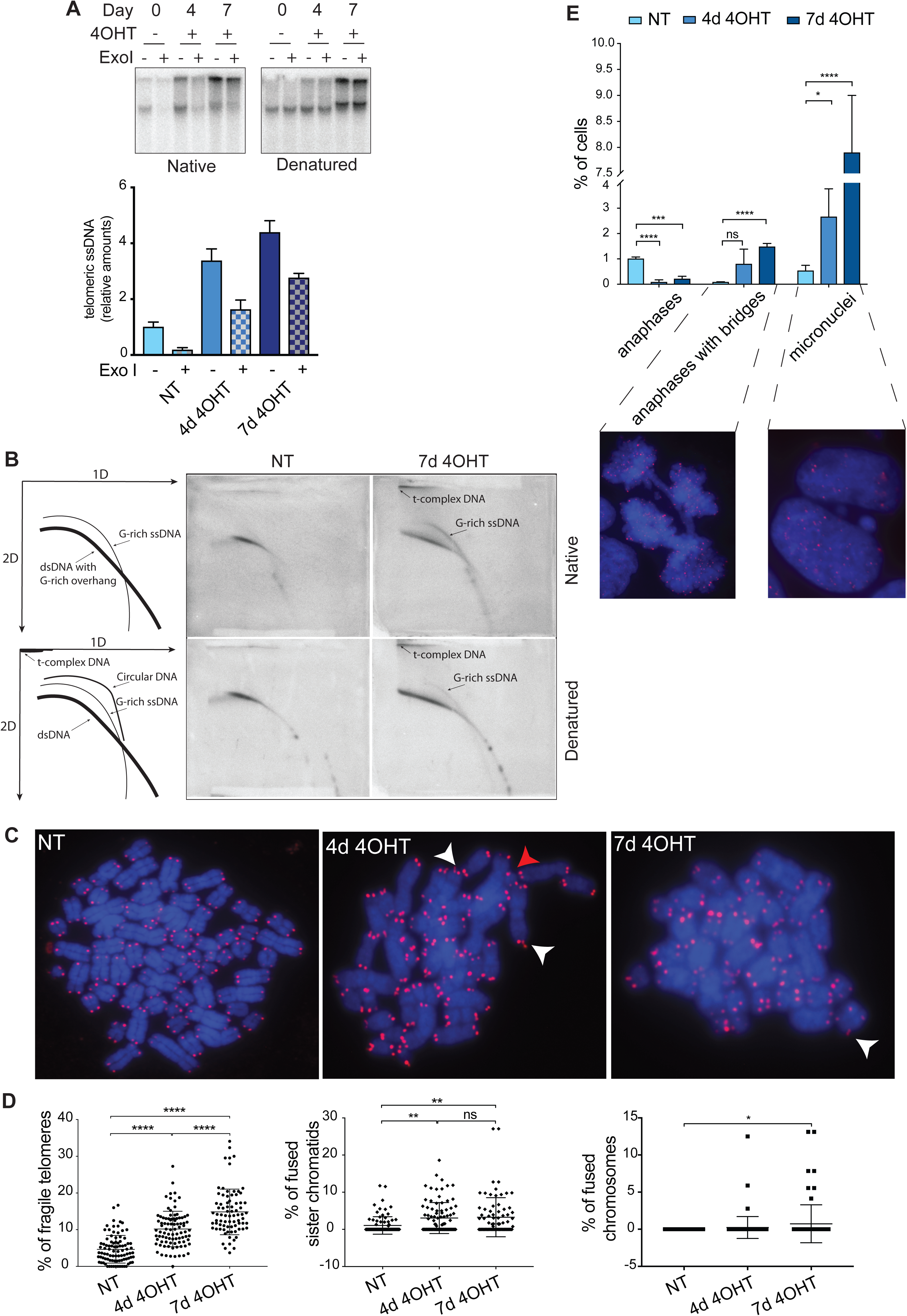
POT1 Loss Leads to Accumulation of Single-Stranded G-Rich DNA, Branched DNA Structures and Telomere Fragility. (A) G-overhang assay for clone 35 upon POT1 removal. The bars show relative amounts of single stranded telomeric DNA + SD. Experiments were performed in triplicate. (B) 2D gels for clone 35 derive DNA with POT1 (NT) and upon POT1 removal for 7 days (7d 4OHT). (C) Examples of metaphase spreads for non-treated (NT) clone 35 cells and clone 35 treated with 4-OHT for 4 and 7 days. White arrowheads indicate fragile telomeres, the red arrowhead indicates a sister-chromatid fusion. (D) Quantification of fragile telomeres (left graph), sister chromatid fusions (middle graph) and chromosome fusions (right graph) as a percentage of events per metaphase. Experiments were performed in triplicate. At least 25 metaphases were examined per condition per replicate. The mean is displayed and error bars represent ± SD. Significance was determined using one-way ANOVA. * p<0.05, ** p<0.01, *** p<0.005, **** p<0.0001. (E) Quantification and representative images of anaphases with bridges and micronuclei for non-treated (NT) clone 35 cells and clone 35 treated with 4-OHT for 4 and 7 days. Experiments were performed in triplicate. At least 650 cells were examined per condition per replicate. Significance was determined using one-way ANOVA. ns – not significant. * p<0.05, ** p<0.01, *** p<0.005, **** p<0.0001. In all experiments, POT1 deletion was induced with 0.5 μM 4-OHT.

To further characterize the telomeric DNA abnormalities we used two-dimensional gel electrophoresis (Wang et al., 2004). DNA was digested with HinfI and RsaI and electrophoresis in the first and second dimension was performed in the absence and presence of ethidium bromide to estimate DNA size and conformation, respectively (Figure 2B). Telomeric DNA was detected before and after denaturation. In wild type cells, a single arc was detected consistent with linear double stranded telomeric DNA with a 3’ overhang (Wang et al., 2004). *POT1* deletion led to the appearance of the slow migrating so-called t-complex (Nabetani and Ishikawa, 2009) which was particularly retarded in the presence of ethidium bromide and which therefore is considered to represent branched DNA with double stranded and single stranded portions (Nabetani and Ishikawa, 2009). In addition, single stranded G-rich DNA appeared as an arc, which may stem from unbranched molecules (Figure 2B).

Telomere replication defects lead to accumulation of so-called fragile telomeres in which the telomeric DNA signal detected on metaphase chromosomes is fuzzy or sometimes split into two (Sfeir et al., 2009). We quantified telomere fragility as well as other telomere abnormalities in *POT1* wild type and knockout cells (Figure 2C and 2D). We observed a strong increase in telomere fragility from 5% to 10% after 4 days and to 15% after 7 days of *POT1*-deletion. Telomere sister chromatid fusions and chromosome end-to-end fusions also increased but this concerned a much smaller fraction of telomeres (Figure 2D). Consistent with the analysis by Southern blotting, the metaphase chromosomes of *POT1* knockout cells showed telomere lengthening and a striking increase in telomere length heterogeneity. When observing interphase nuclei, we noticed increased amounts of micronuclei, which is an indication for genomic instability upon POT1 loss (Figure 2E). Finally, while anaphase cells were rare to detect in *POT1* knockout cells, they displayed increased frequency of anaphase bridges, which were likely caused by the chromatid and chromosome fusions (Figure 2E).

### The POT1 Loss Phenotype Is Caused by Deprotection of the G-rich DNA Overhang

The POT1 protein contains two OB-folds in its N-terminal half with which it binds single stranded telomeric DNA (Lei et al., 2004). The C-terminal half contains a third split OB-fold which is bound by TPP1 via multiple interactions (Chen et al., 2017; Rice et al., 2017). For telomere recruitment, the interaction of POT1 with TPP1 is necessary and sufficient (Liu et al., 2004; Pinzaru et al., 2016; Ye et al., 2004). In order to test if DNA binding by POT1 is required for averting the described telomere abnormalities we developed a complementation system in which triple HA-tagged POT1 was expressed in the *POT1* knockout cells from a lentiviral construct containing a doxycyclin inducible promoter (Figure 3A). Expression of exogenous POT1 rescued the knockout cells from the DNA damage response and the structural abnormalities in telomeric DNA (Figures 3A and 3B). Expression of mutant POT1 versions, some of which are associated with cancer development and which in previous analyses showed reduced but not abolished single stranded DNA binding, could also avert the severe telomere abnormalities when expressed from the plasmid (Figure 3C, 3D and S3A). However, two POT1 mutants were unable to rescue the telomere defects. The POT1F62V point mutation which diminishes single strand telomeric DNA binding affinity by a factor of 65 (Pinzaru et al., 2016), abolished POT1 function with regard to telomeric DNA structure (Figure 3E) as well as DNA damage response (Figure S3B). The same was true for POT1ΔOB lacking the N-terminal OB fold. Therefore, we conclude that the direct binding of single stranded telomeric DNA by POT1 is required to prevent telomeric DNA abnormalities. The telomere recruitment via TPP1 is not sufficient for POT1 function.

**Figure 3.**
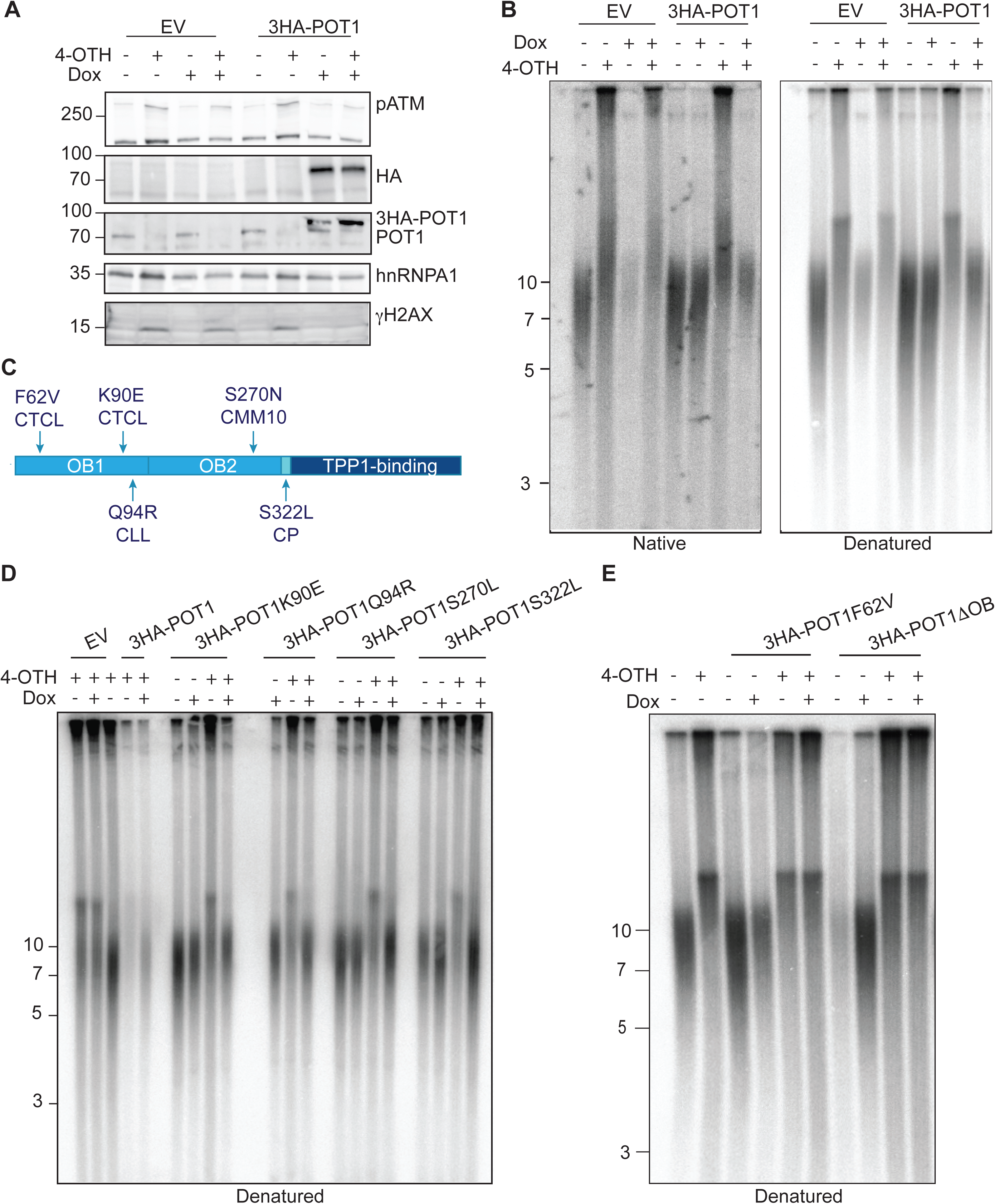
The POT1 Loss Phenotype Is Caused by Deprotection of the G-rich DNA Overhang. (A) Western blot for POT1 and DDR markers upon POT1 removal in clone 35 and overexpression of ectopic WT POT1. (B) Telomere length and G-overhang length analysis for clone 35 upon POT1 removal and overexpression of ectopic WT POT1. (C) Schematic of diseases-associated mutations in POT1. CLCL - cutaneous T cell lymphoma, CMM - cutaneous malignant melanoma, CLL - chronic lymphatic leukemia. (D), (E) Telomere length and G-overhang length analysis for clone 35 upon POT1 removal and overexpression of ectopic POT1 variants carrying disease-associated mutations. In all experiments, POT1 deletion was induced with 0.5 μM 4-OHT for 7 days. Expression of ectopic POT1 was induced with 1 μg/ml doxycycline for 4 days.

As our knockout system did not affect a shorter polypeptide derived from the *POT1* locus referred to as POT1-55 ((Hockemeyer et al., 2005); Figure 1A), we used lentiviral-mediated delivery of shRNA that targets both POT1 forms (Hockemeyer et al., 2005) to confirm that the phenotype we observed was POT1-55 independent (Figure S3C). Indeed, the telomere elongation and DNA damage response phenotypes upon *POT1* knockout induced with 4OHT and POT1 knockdown with shRNA were identical (Figure S3D).

### Development of 2-Step QTIP Uncovers Substantial Remodeling of the Telomeric Proteome Upon POT1 Loss

We expected that the above described telomere abnormalities in the absence of POT1 would trigger substantial changes in the telomeric protein composition. In addition, we considered the possibility that the telomeric DNA abnormalities in *POT1* knockout cells might be caused by inappropriate association of DNA processing enzymes with telomeric DNA in the first place. In order to address these questions, we embarked on isolating and analyzing the telomeric proteome in presence and absence of POT1. Our lab previously developed a quantitative telomeric chromatin isolation protocol (QTIP) in which crosslinked telomeric chromatin is purified with antibodies against the abundant telomere binding proteins TRF1 and TRF2 and analyzed by mass spectrometry (Grolimund et al., 2013; Majerská et al., 2017). Different telomeric states were compared by differential SILAC labeling. Here, we wanted to compare telomeric protein composition in a larger set of different samples and therefore opted for the use of tandem mass tag reagents (TMT) (Hogrebe et al., 2018; O’Connell et al., 2018; Rauniyar and Yates, 2014). In addition, we added to the QTIP method an additional affinity purification step by first purifying the crosslinked and sonicated telomeric chromatin with anti-FLAG sepharose beads containing monoclonal antibodies against the FLAG epitope (Figure 4A). This step became possible as our cell line contained FLAG-tagged TRF1 and TRF2. Immunoprecipitated chromatin was eluted with excess of FLAG-peptide and then re-precipitated with polyclonal affinity-purified anti-TRF1 and anti-TRF2 antibodies as in the classical QTIP method. We refer to this method as 2-step QTIP. The two consecutive affinity purification steps yielded a recovery of telomeric DNA of 38-47% and an up to more than 1,000x fold enrichment of telomeric DNA over Alu-repeat DNA (Figure 4B).

**Figure 4.**
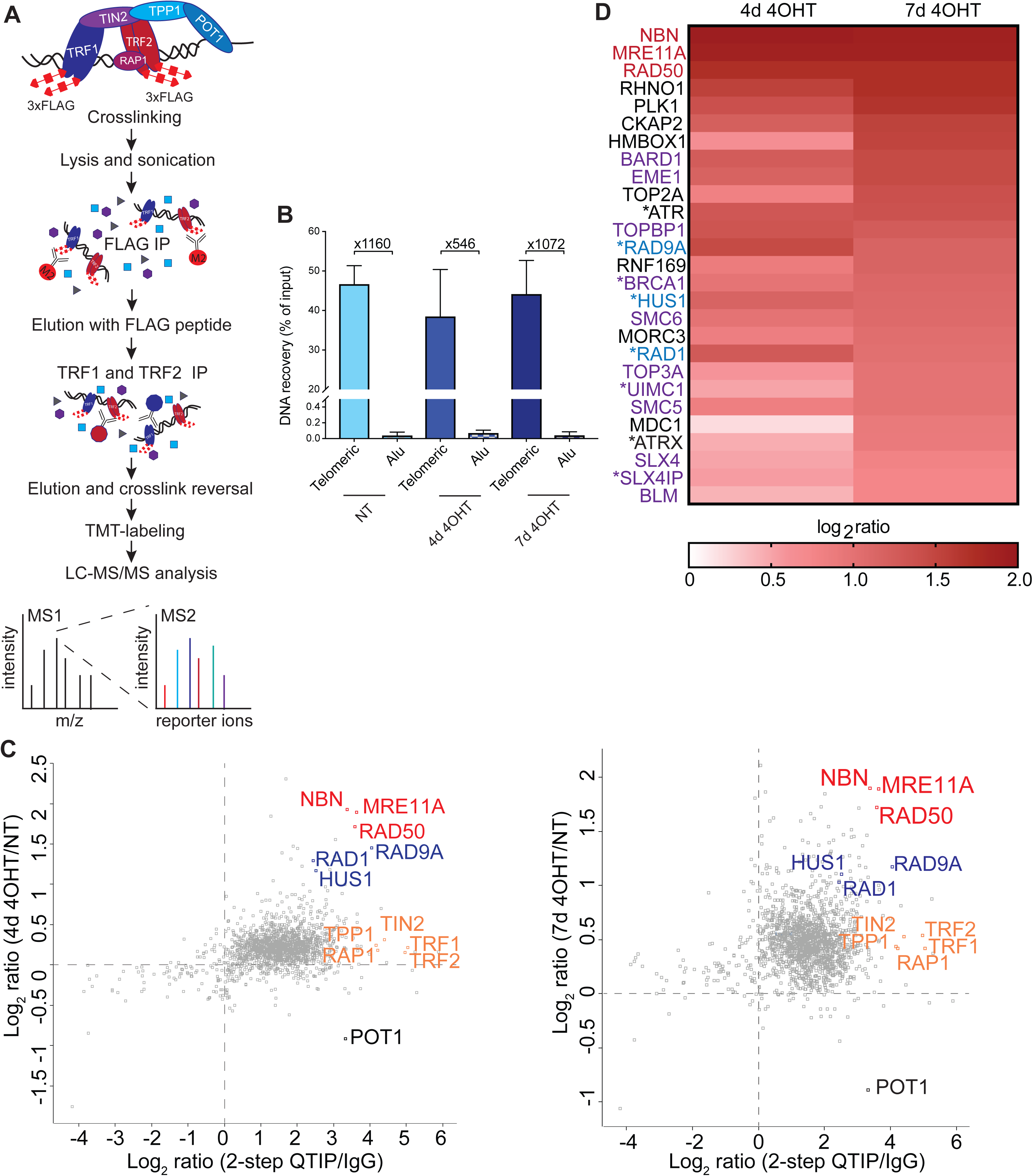
Development of 2-Step QTIP Uncovers Substantial Remodeling of the Telomeric Proteome Upon POT1 Loss. (A) Schematic drawing of 2-step QTIP (improved version of the quantitative telomeric chromatin isolation protocol (QTIP)). (B) Quantification of precipitated telomeric DNA in 2-step QTIP (% of input) and fold enrichment of precipitated telomeric DNA compared with precipitated Alu repeat DNA (based on dot blot analyses). Bars represent data from three independent experiments + SD. (C) Scatter plots representing immunoprecipitation specificity (2-step QTIP/IgG ratio) versus difference between non treated cells (NT) and cells at day 4 upon 4-OHT induction (4d 4OHT/NT ratio) or difference between non treated cells (NT) and cells at day 7 upon 4-OHT induction (7d 4OHT/NT ratio) respectively (data from Supplementary Table 1). Values are the average of the 3 biological replicates. (D) Heat map representing changes in levels of selected proteins involved in DNA damage (MRN complex in red, 9-1-1 complex in blue) and homology recombination (purple) after *POT1* knockout induction. Proteins marked with * pass t-test, but not t-test with Benjamini-Hochberg correction (p<0.05, q<0.09).

We performed three biological replicates with parental cells and cells from which *POT1* had been deleted for 4 or 7 days (Figure 4C). To increase throughput and reduce the analytical variability and the incidence of missing data, we applied a 6-plex TMT isobaric labelling workflow to analyze 2-step QTIP preparations of untreated (NT) and *POT1* knockout cells 4 and 7 days post induced deletion. In addition, we included IgG negative controls. Each of the three experimental replicates thus constituted a distinct TMT mix. After digestion, labeling and pooling, peptides in each mix were extensively fractionated by peptide isoelectric focusing to maximize separation and increase depth of protein identification. Spike-in proteins were used to monitor and correct signal levels across replicates and assess the dynamic range of TMT quantitation, which is known to display ratio compression phenomena (Savitski et al., 2013) (Figure S4A and S4B). To maximize sensitivity, we chose not to apply synchronous precursor selection scan methods (McAlister et al., 2014). Analysis by mass spectrometry and comparison of 2-step QTIP purified chromatin to IgG control purified chromatin indicated that the 2-step QTIP approach yielded telomeric chromatin of unprecedented purity. Indeed, the vast majority of proteins detected were clearly enriched in the QTIP samples compared to the IgG controls (Figure 4C, Supplementary Table 1). The 2-step QTIP procedure with fractionation allowed to detect, in addition to the most abundant shelterin components, more than 1400 proteins. When comparing the telomeric chromatin in wild type to *POT1*-knockout cells we observed a slight increase in all shelterin components except POT1, which as expected was strongly diminished (Figure 4C). The slight increase of five of the six shelterin proteins can be explained by the telomere elongation that occurred during the experiment (Figure 1D and 2C). 157 proteins were significantly enriched at telomeres that lacked POT1 at day 7 after knockout induction (paired T-test with Benjamini-Hochberg FDR correction and threshold at 0.05 on the adjusted p-value). Among them, a large set of proteins involved in the DNA damage response and homology directed DNA repair was identified (Figure 4D).

### POT1 Prevents Telomeric Accumulation of Homology-Directed and other Repair Factors, Nucleases and Cell Cycle Regulators

We grouped 88 proteins that were significantly enriched at telomeres in *POT1* knockout cells more than 1.5 fold (log2>0.6) into functional classes using the STRING database (Szklarczyk et al., 2019). The cutoff chosen takes into consideration TMT ratio compression effects (Figure S4). Apart from known telomere maintenance components, we identified checkpoint response factors, nucleases, DNA repair proteins, and proteins involved in cell cycle regulation, chromosome organization, centromere function and RNA metabolism (Figure 5). Interestingly, the largest cluster (15 proteins) contained cell cycle and mitosis-associated proteins (Figure 5A). We identified multiple proteins playing roles in spindle formation (AURKA, INCENP, KIFC1, TPX2, AURKB, KIF20A, PLK1). In the same cluster fall two condensin subunits (NCAPD2 and NCAPG) as well as cyclin B1 (CCNB1) and origin recognition complex subunit 1 (ORC1). In the second largest cluster (11 proteins, Figure 5B) we found DNA damage response and repair proteins as well as HR proteins: all components of MRE11/RAD50/NBS1 (MRN) complex, TOPBP1, RHNO1, BARD1, BLM, TOP3A, and the structure specific endonucleases ERCC1, ERCC4 and EME1. SLX4 and SLX1A which function as Holliday junction resolvases are also linked to this cluster. Another cluster included proteins involved in RNA metabolism (9 proteins, Figure 5C), which may relate to the increased abundance of R-loops at POT1-depleted telomeres (see below). We also identified a cluster of proteins that contained subunits of the SMC5-SMC6 complex, which is involved in HR as well as TOP1, SUMO1 and SUMO2 (Figure 5D). The latter might stem from sumoylated proteins present at damaged telomeres. Also among proteins enriched at telomeres lacking POT1 we identified SMCHD1 and LRIF1 that have been previously implicated in the telomeric DNA damage response (Vančevska et al., 2019), parts of the protein phosphatase 1 complex and lamin A that had been identified at telomeres previously (Kim et al., 2009; Briod et al., accompanying paper).

**Figure 5.**
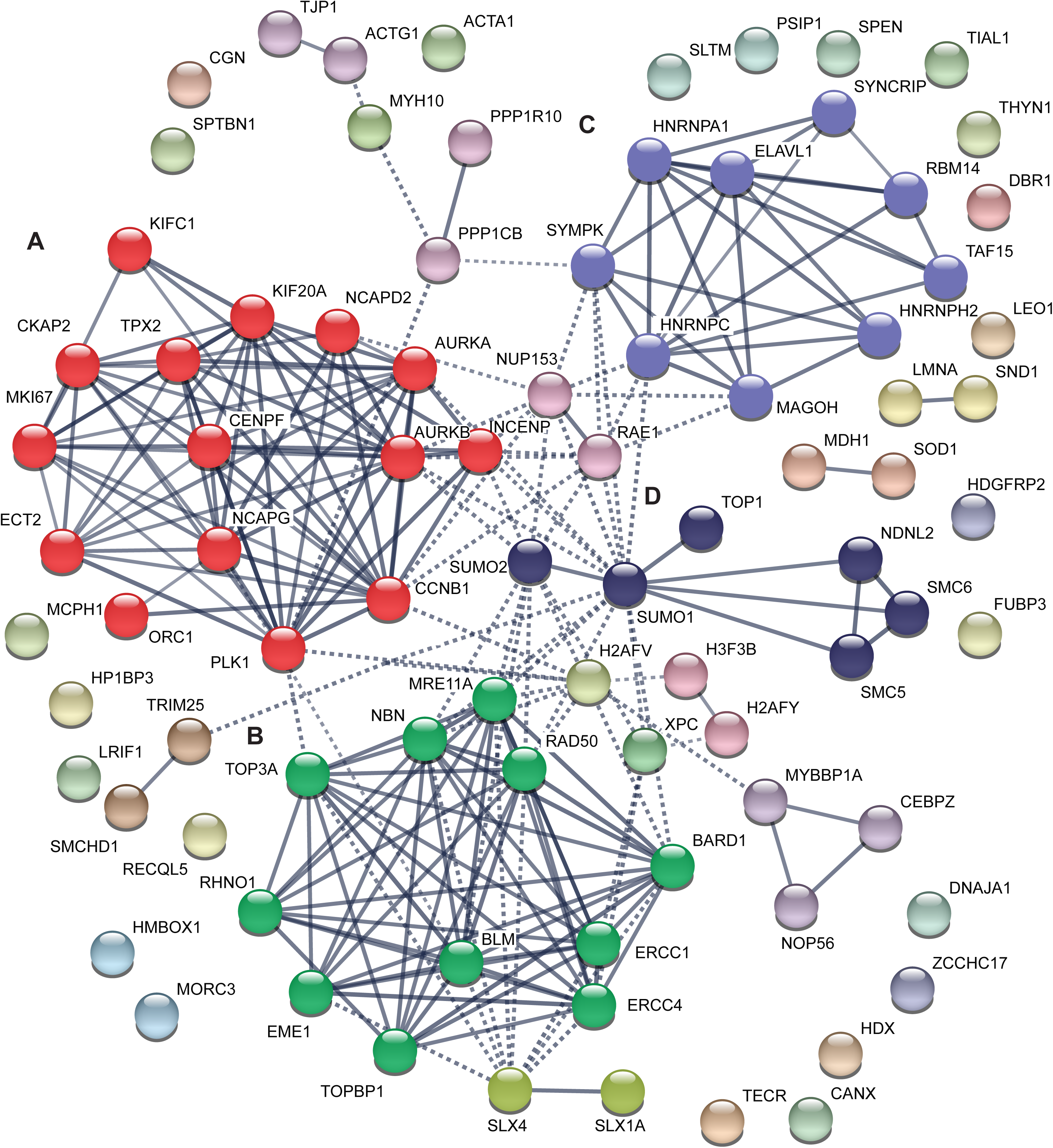
POT1 Prevents Telomeric Accumulation of HDR and other Repair Factors, Nucleases and Cell Cycle Regulators. Cluster analysis of proteins enriched at telomeres lacking POT1 using STRING. The largest identified clusters are colored in red (mitosis-associated proteins, (A)), green (recombination and DNA damage response proteins, (B)), purple (RNA-binding proteins, (C)), and dark blue (chromatin and DNA remodelers, (D)). Line thickness indicates the strength of data support.

### Telomere Elongation and Branching upon POT1 Loss is Mediated by the HDR Machinery

We next embarked on testing if the identified proteins played roles in establishing or responding to the POT1 loss mediated phenotype. Since POT1 loss caused telomere elongation, we were particularly intrigued by the enrichment of the HDR factors MRE11, BARD1, BRCA1 and BLM at telomeres. Very strikingly, when we depleted these and other HR factors (RAD51, BRCA1, BRCA2, BARD1, BLM) by siRNA in *POT1*-knockout cells (Figures S5A, S5B, S5C and S5D), telomere elongation, telomere branching and G-rich single strand DNA accumulation were all suppressed (Figure 6A and 6B). These data therefore indicate that in the absence of POT1 the HDR machinery becomes overly active at telomeres causing severe telomere instability. As POT1 was previously implicated in regulation of telomerase (Loayza and De Lange, 2003; Kelleher et al., 2005; Latrick and Cech, 2010; Zhong et al., 2012) we also tested, if telomerase may contribute to the POT1 loss mediated phenotype adding the telomerase inhibitor GRN163L to cells at the same time when *POT1* deletion was induced (Figure 6C). However, although GRN163L strongly reduced telomerase activity (Figure S5E), the POT1-loss mediated phenotype remained unaltered. Thus, telomerase showed no contribution to the phenotype in this experiment. Finally, we tested if HDR inhibition influenced the DDR observed upon POT1 loss by detecting pCHK1 or γ-H2AX on Western blots (Figure S5A and S5B). However, in *POT1* knockout cells the DDR still occurred upon suppression of HDR.

**Figure 6.**
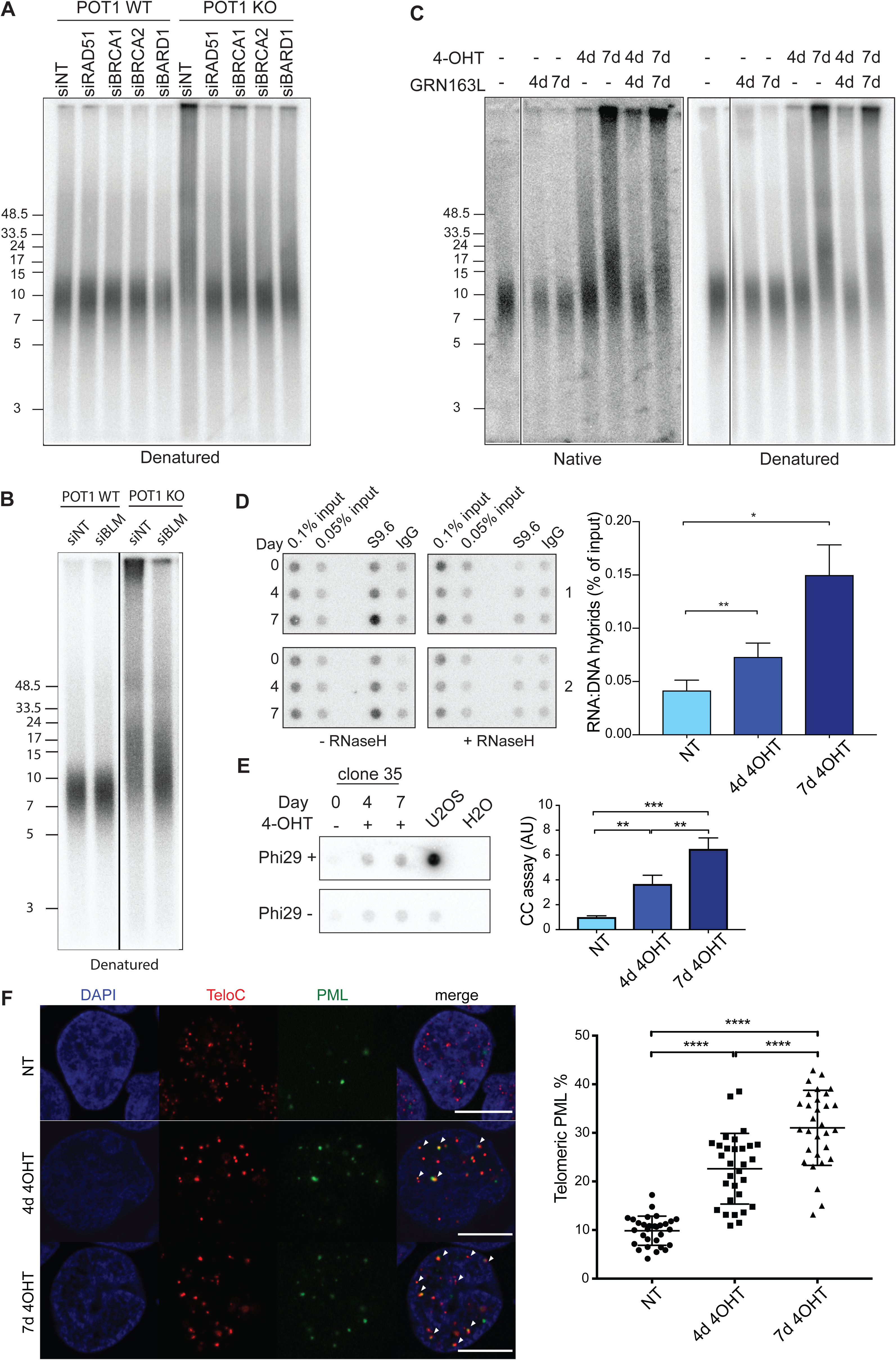
Telomere Elongation and Branching under POT1 Loss is Mediated by the HDR Machinery. (A), (B) Telomere length for clone 35 upon POT1 removal and depletion of selected recombination proteins with siRNA. (C) Telomere length and G-overhang length analysis for clone 35 upon POT1 removal and inhibition of telomerase with 1µM GRN163L. (D) DRIP assay for DNA-RNA hybrid accumulation at telomeres in clone 35 upon POT1 removal. Dot blots represent two replicates. Bars represent data from three independent experiments +SD. The IgG signal was subtracted. Significance was determined using one-way ANOVA, * p<0.05, ** p<0.01. (E) C-circle (CC) assay for clone 35 upon POT1 removal. Bars represent data from three independent experiments +SD. Significance was determined using one-way ANOVA, ** p<0.01, *** p<0.005. In all experiments, *POT1* deletions was induced with 0.5 μM 4-OHT for 4 and 7 days. (F) Quantification and representative pictures for visualization of the PML protein (IF, green) and telomeres (FISH, red) in non-treated (NT) clone 35 cells and clone 35 treated with 4-OHT for 4 and 7 days. Arrows indicate co-localizations. Scale bar equals 10 µm. Experiments were performed in triplicate. At least 150 cells were examined per condition per replicate. Significance was determined using one-way ANOVA. **** p<0.0001. In all experiments, POT1 deletion was induced with 0.5 μM 4-OHT.

### POT1 Loss Leads to Accumulation of Telomeric R-loops, C-circles and Telomeric PML Bodies

Next we assessed for presence of telomeric DNA-RNA hybrid structures which can be formed with the telomeric long noncoding RNA TERRA and are suppressed at telomeres in wild type cells by RNA surveillance factors (Azzalin et al., 2007; Chawla et al., 2011), RNase H (Arora et al., 2014; Graf et al., 2017), FANCM (Silva et al., 2019) and the THO-complex (Pfeiffer et al., 2013) but can accumulate in mutant conditions interfering with telomere replication. Strikingly, telomeric R-loops are a hallmark of ALT cells which maintain their telomeres by DNA recombination (Arora et al., 2014; Graf et al., 2017). To detect telomeric R-loops, the structure specific antibody S9.6 was used to precipitate from total nucleic acid the R-loops. Telomeric signals in precipitated nucleic acid were quantified by dotblot hybridization (Figure 6D). As a control for the specificity for R-loop precipitation, the nucleic acid samples were treated in vitro prior to immunoprecipitation with commercial RNase H. RNase H destroys the RNA moiety in DNA/RNA hybrids. As expected, in vitro RNase H treatment abolished the telomeric signal to background levels validating the specificity of the immunoprecipitation for R-loops (Figure 6D). *POT1* deletion caused increased accumulation of telomeric R-loops in the time course experiment. At the same time, *POT1* deletion did not significantly affect TERRA expression levels as detected on Northern blots (Figure S6A) and it did not significantly affect individual TERRA molecules when the quantities stemming from six different chromosome ends were analyzed by RT-qPCR (Figure S6B).

Another hallmark of ALT cells is the formation of so-called C-circles which are single stranded circular DNA molecules containing telomeric C-strand repeats of unknown function (Henson et al., 2017). They can be detected by rolling circle amplification followed by hybridization. *POT1* deletion caused a significant increase of C-circles, which however was lower than then the amounts detected in U2OS ALT-cells (Figure 6E).

Finally, we examined the control cells and *POT1* knockout cells for presence of promyelotic leukemia (PML) bodies, which represent another marker for ALT activity (Yeager et al., 1999). The percentage of PML foci that were present at telomeres increased upon *POT1* deletion significantly from 10 to more than 30% after 7 days of induced deletion (Figure 6F). Altogether, with increased telomeric R-loops, C-circles and PML bodies the POT1 loss promoted the establishment of three markers that are typical for ALT cells.

## DISCUSSION

### Conditional Deletion of POT1 Uncovers Major Roles in Telomere Stability

In this paper we uncover crucial roles of human POT1 in repressing HDR at telomeres which gives rise to severe telomeric DNA abnormalities including telomere elongation, telomere branching, accumulation of single stranded telomeric DNA and telomeric R-loops. The establishment of a conditional system as well as the application of telomere proteomics were crucial to obtain the documented insights. POT1 function had been studied before in several systems but despite its conservation in evolution and the conserved association with G-rich single stranded telomeric DNA, POT1 loss gave different phenotypes in different model systems. From the work in *S. pombe*, POT1 appeared to be most important for the protection of chromosome ends from nucleolytic degradation (Baumann and Cech, 2001). In mice and other rodents, the *POT1* gene is duplicated and different functions have been assigned to the two paralogs indicating remarkable divergence from humans in telomere biology. With mouse cells, the data about a striking function of POT1 paralogs in the suppression of HDR were controversial (Hockemeyer et al., 2006; Wu et al., 2006), although the data seemed to support subtle contributions in the suppression of HDR at least in mutant backgrounds lacking the Ku end binding protein (Palm et al., 2009). In previous depletion studies of human POT1 (Hockemeyer et al., 2005; Kim et al., 2017), low amounts of POT1 may have been retained being sufficient to efficiently suppress telomeric HDR. Residual amounts of POT1 in RNA interference experiments may also explain why the telomeric damage that had been detected by Hockemeyer et al. (2005) was only prominent in G1, and not in S and G2 phases of the cell cycle, whereas we see DNA damage at telomeres throughout cell cycle in our system. From this one may conclude that absence of POT1 can be tolerated in G1 phase of cell cycle in immortalized cells, and that low levels of POT1 are sufficient at telomeres in S and G2 cells to suppress telomeric HDR that leads to severe telomere instability, enhanced DNA damage response and possibly, mitotic catastrophe. Also consistent with this notion, we found that POT1 mutants that had been detected in cancer were able to suppress HDR despite the fact that their binding ability of DNA was reduced.

We also observed a significant increase in telomere fragility in the cells lacking POT1 during the time course experiment (Figure 2B). Telomere fragility is a hallmark of replication stress (Sfeir et al., 2009) and in ALT cells telomere fragility has been suggested to be caused by DNA recombination events among telomeric sequences (Min et al., 2017) being consistent with our findings. Interestingly, disease-associated POT1 mutations have been shown to increase telomere fragility and cause replication fork collapse when overexpressed (Pinzaru et al., 2016). We suspect that these effects may have been due to slightly increased recombination events at telomeres due to POT1 dysfunction, which may have led to accumulation of recombination by-products and R-loops at telomeres that further challenged replication of telomeric sequences.

In previous work it has been demonstrated that POT1 can regulate telomerase either negatively or positively (see Introduction). However, we observe that complete absence of human POT1 in HEK293E cells leads to rapid telomere elongation that depends on HDR and does not require telomerase (Figure 6C). Thus, our data reveal that POT1 plays crucial roles not only in regulating telomerase but also in suppressing recombination-based mechanisms of telomere elongation. Overall, we suspect that the divergence of observed phenotypes upon POT1 depletion or *POT1* knockout may not reflect fundamental differences in telomere function of POT1 in different model systems. We rather imagine that system-specific variabilities with regard to functional redundancy and abundance of various telomeric proteins may strongly influence the phenotypic outcomes. Whereas POT1 emerges as a master in the protection and regulation of single strand telomeric DNA, in its absence RPA, CST, RAD51, hnRNPA1, RADX and telomerase may all compete for binding to the G-rich single stranded telomeric DNA and guide the faith of chromosome ends towards different outcomes.

### 2-Step QTIP and TMT Labeling Unravels the Telomeric Proteome

We developed and implemented here a 2-step QTIP protocol in order to obtain telomeric chromatin of high purity and TMT labeling in order to quantitatively compare the telomeric proteome changes obtained upon POT1 loss. TMT labeling yields highest precision but it suffers from a ratio compression phenomenon and thus underestimates differences in protein abundance (Hogrebe et al., 2018; O’Connell et al., 2018; Rauniyar and Yates, 2014). Nevertheless, we could identify a large set of proteins whose association with telomeres is suppressed by POT1. However, due to variability between biological replicates several DNA damage proteins that we used as positive controls (for example RAD9A-HUS1-RAD1), did not pass our stringent threshold (paired T-test with Benjamini-Hochberg FDR correction and threshold at 0.05 on the adjusted p-value) when considering individual subunits. Therefore, we also considered candidates with unadjusted p-values <0.05, especially when all subunits of multiprotein complexes behaved in a consistent manner. As expected for the ATR-dependent DNA damage response, the 9-1-1 complex components and TOPBP1 were enriched at telomeres already at day 4 post induction. Their abundance decreased slightly at day 7 when recombination events became most prevalent. All the other identified DNA damage and HDR proteins were most enriched at telomeres 7 days after induction. Among the 157 proteins that were significantly enriched, 88 were enriched more than 1.5 fold (log2>0.6) (Figure 5). Apart from the DNA damage proteins whose association was expected, we identified a number of functional protein groups that could not be anticipated. The increased presence of proteins involved in RNA metabolism at POT1-depleted telomeres may be related to the increased amounts of R-loops at telomeres lacking POT1 (Figure 6D). The observed enrichment of mitosis and spindle-associated proteins may stem from the perturbed mitosis in cells lacking POT1 (Figure 2E). The functional relevance of HDR proteins was demonstrated in the current work while for others their exact roles remain to be defined.

### Human POT1 Suppresses Damage-Inducing HDR

Our work supports the following model for human POT1 function (Figure 7). In wild type cells, the single stranded telomeric DNA is to a large extent covered by POT1. Upon POT1 loss, the telomeric C-strand is further resected and RPA binds, triggering an ATR dependent checkpoint response. Eventually, RPA is at least partially replaced by RAD51 presumably through BRCA2-mediated loading as during classical recombination. RAD51-coated telomeric 3’ overhangs will invade telomeric DNA of adjacent chromosome ends, which then will be extended by DNA polymerases. The strand invasion is responsible for the observed branched DNA structures. The observed extended 3’ overhangs are due to 3’ end extension upon strand invasion and the presumed 5’ end resection, which follows POT1 loss. In addition, the increased abundance of R-loops between TERRA and the telomeric C-strand should increase single stranded G-rich telomeric DNA and may facilitate RAD51 binding and telomere recombination (Arora et al., 2014; Graf et al., 2017). The observed telomere fragility is due to abnormal chromatin condensation following HDR or gaps in telomeric DNA that may remain after the repair process. Overall, our data illustrate that while homology directed repair is highly beneficial to safeguard the genome from chromosomal breaks, it becomes detrimental when out of check at the natural ends of chromosomes, leading to severe telomere and chromosomal instability. The POT1 protein masters the repression of HDR through its sequence specific DNA binding, covering the single stranded telomeric DNA. POT1 therefore prevents the unwanted engagement of the HDR repair machinery at telomeres but not elsewhere in the genome.

**Figure 7.**
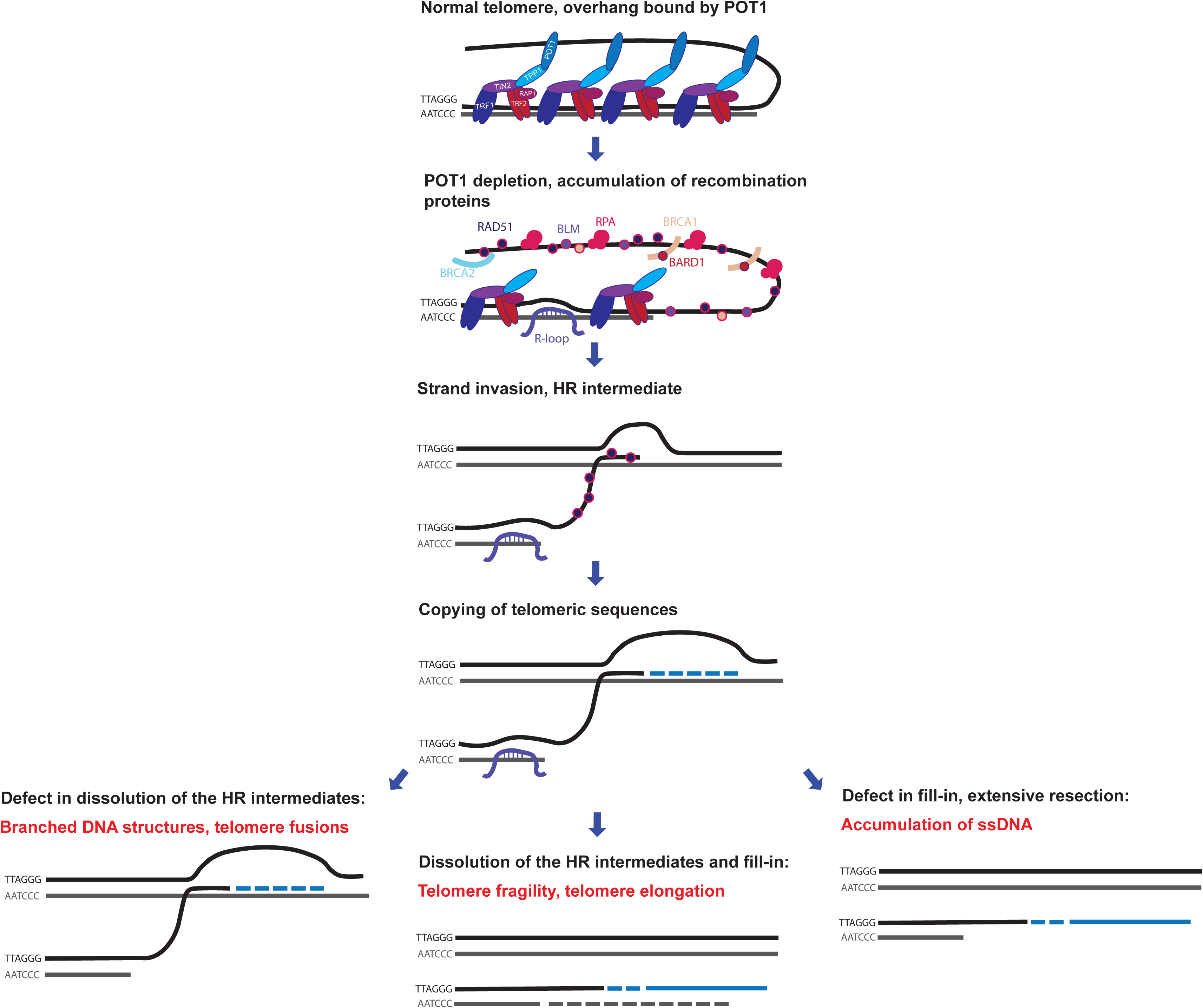
Human POT1 Suppresses Damage-Inducing HDR. In the absence of POT1 the telomeric C-strand is resected and RPA and recombination proteins associate with the overhang. RAD51-coated overhangs invade telomeric DNA of adjacent chromosome ends and the invading 3’end is extended by DNA polymerases. Accumulation of R-loops at telomeres facilitates strand invasion. In case of proper dissolution of the HR intermediates telomeres are elongated and fragility is due to abnormal chromatin condensation after HDR. In case of defective dissolution of the HR intermediates leads to accumulation of branched DNA structures. Incomplete fill-in synthesis leads to accumulation of G-rich single stranded DNA.

## SUPPLEMENTAL INFORMATION

Supplemental Information includes 6 figures and 2 Supplementary Tables and can be found with this article online.

## ACKNOWLEDGEMENTS

We thank members of the Lingner lab for fruitful discussions, the Protein Expression Core Facility at EPFL for 293E cells, Gérald Lossaint for 293E cl75, Pierre Gönzy for U2OS cells, Etienne Meylan for the pCW22 construct and Jerry Shay (UT Southwestern) for GRN163L. Research in J.L.’s laboratory was supported by the Swiss National Science Foundation (SNSF), the SNSF funded NCCR RNA and disease network, an Initial Training Network (ITN) grant (aDDRess) from the European Commission’s Seventh Framework Programme, the Swiss Cancer League and EPFL.

## AUTHOR CONTRIBUTIONS

GG carried out all experiments except for the mass spectrometry analysis, which was done by MQ. AB carried out initial experiments demonstrating that RAD51 depletion rescues from telomeric DNA abnormalities observed in *POT1* knockout cells. GG and JL wrote the paper. The study was conceptualized by GG and JL.

## METHOD DETAILS

### Cell lines, CRISPR/Cas9 gene editing, transfections and lentivirus production

HEK293E cells were grown in DMEM with 10%FCS and penicillin/streptomycin solution (1:100) at 37°C in 5% CO_2_. The population doubling (PDL) values were calculated using the mathematical formula PD = [(ln(N2)) - (ln(N1))] / ln(2) where N1 is the number of cells plated and N2 the number of cells harvested.

293E cl75p100 clone was generated from 293E cl75 by transfection of pSpCas9(BB)-2A-GFP (PX458) plasmid expressing gRNA targeting exon 6 of *POT1* (gRNA1, 5’-CACCGCCCCTGAATCAACTTAAGGG-3’), a recombination reporter plasmid containing a binding site for gRNA1 (pMB1610_pRR-Puro) and pUC19 plasmid containing a repair template for exon 6 of *POT1* (with a modified gRNA binding site) flanked with LoxP sites and homology arms of 500 bp (pUC19_RT). Cells were selected with 1 µg/mL puromycin for 5 days and clones were isolated by single cell dilution.

Clones were screened using 2 genotyping PCRs (with OneTaq polymerase premix from NEB and 1.25 µM primers 1+2 and 1+3). Positive clones were kept in culture for several weeks to monitor telomere length stability and POT1 levels. The gRNA was designed using http://crispr.mit.edu/ and cloned into pSpCas9(BB)-2A-GFP (PX458).

Cre clones 35 and 45 were generated by transducing cl75p100 cells with lentivirus containing CreERT2, selecting cells with 1 µg/mL puromycin for 5 days and isolation of clones by single cell dilution. Isolated clones were tested for *POT1* deletion by inducing Cre recombinase with 0.5 µM 4-OHT and clones with the highest recombination efficiency were kept and tested for telomere length stability. Cells were transfected with plasmid DNA using Lipofectamine 2000 (1:200) and 2.5 µg of DNA in 2 mL of OPTIMEM per well in 6-well plates. One day after transfection, cells were split and selected with either puromycin (1 µg/mL) or blasticidin (10 µg/mL). Cells were harvested or diluted 5 days after transfection.

Cells were transfected with siRNA pools from Dharmacon using calcium-phosphate transfection with 10 pmol of siRNA in 1 mL of DMEM without antibiotics per well in 6 well plates. One day after transfection, cells were split and 72 hours after transfection cells were harvested for TRF analysis, q-RT PCR and Western blot analysis. For lentiviral production 293T cells were transfected in 10 cm dishes with 4 µg of the corresponding lentiviral vectors (shRNA or transgene) and packaging vectors pMD2.G (1 µg) and pCMVR8.74 (3 µg) using Lipofectamine 2000 (1:400 in OPTIMEM). The next day, the medium was changed for DMEM with 10% FCS and penicillin/streptomycin. The first harvest of viral particles was done 40 hours after transfection and fresh medium was added to the cells. Virus-containing medium was filtered through 0.45 µm filters and 1 ml of filtrate was applied to 1 million of cells in one well of a 6-well plate. The second harvest was done 24 hours after the first harvest and the transduction was repeated as described. The next day transduced cells were split and selected with puromycin (1 µg/mL), blasticidin (10 µg/mL) or hygromycin (100 µg/mL) for at least one week. For doxycycline-inducible expression of POT1, HA-tagged WT POT1 or mutant POT1 cDNAs were cloned into pCW22 lentiviral vector between HpaI and PacI restriction sites. Stable cell lines were created by lentiviral transduction and blasticidin selection. Ectopic expression of POT1 constructs was induced by 1µg/mL doxycycline. For ectopic expression of inducible Cre recombinase ERT2CreERT2 was subcloned into pLKO1_puro digested with PpuMI.

### Western blots

1 million cells were resuspended in 100 µl of Laemmli buffer (150 mM Tris-HCl pH 6.8, 4% SDS, 5% beta-mercaptoethanol, 20% glycerol, bromophenol blue) and were boiled for 5 minutes at 95°C. Equivalents of 100,000 cells were loaded on Mini-PROTEAN TGX™ 4-15% acrylamide gels. Primary antibodies were anti-ATM pS1981 (1:1,000, ab81292; Abcam), anti-γH2AX (1:1,000, 05-636; Millipore), anti-Chk1-pS345 (1:1000, #2348, Cell signaling), anti-Chk1 (1:1000, sc-8408, Santa-Cruz), anti-RPA32 –S33 (1:1000, A300-246A, Bethyl), anti-vinculin (1:3000, ab129002, Abcam) and anti-HA (1:3000, BLG-901502, BioLegend). Secondary antibodies were HRP-conjugated goat anti-mouse (W4021, Promega) and goat anti-rabbit (W4011, Promega). For Western blot analysis of POT1, the membrane was denatured after transfer and re-natured with decreasing concentrations of guanidine-HCl (Wu et al., 2007), blocked with 5% milk in TBST and incubated with anti-POT1 (1:1000, ab124784, Abcam) in 5% milk overnight.

### Telomere restriction fragment analysis and G-overhang assay

Genomic DNA from 3-5 million cells was extracted with phenol-chloroform-isoamyl alcohol (25:24:1). 5-10 µg of genomic DNA were digested with 50U HinfI and 30U RsaI in CutSmart buffer (NEB) overnight at 37°C. 3 µg of digested genomic DNA per lane was resolved by pulse field gel electrophoresis in 1% agarose in 0.5xTBE. Samples were electrophoresed for 16 hours at 14°C and 5.2V with 0.5 seconds initial switch and 6 seconds final switch using a CHEF-DRII apparatus (BioRad). The gels were dried for 3 hours at 50°C, prehybridized with Church buffer (0.5 M Na2HPO4 pH 7.2, 1 mM EDTA, 7% SDS, and 1% BSA) and hybridized with a ^32^P-radiolabelled telomeric probe under native conditions (for G-overhang assay) overnight at 50°C. After hybridization, the gels were washed twice for 30 minutes with 2xSSC, 0.5%SDS, 1xSSC, 0.25% SDS, 0.5xSSC, 0.1% SDS and 0.5xSSC, and exposed to a phosphorimager screen. Afterwards the gel was denatured (0.5 M NaOH, 1.5M NaCl), neutralized (0.5 M Tris pH 8.0, 1.5 M NaCl) and re-hybridized as described above. Radioactive signals were detected with Amersham typhoon and quantified in AIDA software version 4.06.034. PRISM7 software was used to calculate relative G-overhang amounts, averages and p-values.

### Two dimensional neutral-neutral agarose gel electrophoresis

Two-dimensional gel electrophoresis was done as described (Wang et al., 2004) with modifications. Equal amounts of HinfI+RsaI digested DNA (10–15 μg) were subjected to electrophoresis on 0.4% agarose gels at 30 V for 16 hours in the dark at room temperature (first dimension). Then, the lanes were cut and overlaid on top of a second gel (second dimension) containing 1.2% agarose and 0.3 μg/mL ethidium bromide. This gel was run at 4°C and 150 V for 6 hours. The gels were processed as described above for the TRF analysis.

### Northern blot

10 µg of RNA per lane was loaded on a 1.2% formaldehyde agarose gel and separated by electrophoresis at 100V for 3 hours. RNA was transferred to a nylon membrane (Hybond N+) overnight in 1X TBE at 15V. The membrane was UV-crosslinked, prehybridized for 1 hour in Church buffer and hybridized with a ^32^P-radiolabeled telomeric probe overnight at 55°C. The membrane was washed in 1xSSC, 0.5% SDS twice for 30 minutes at 60°C and exposed to a phosphorimager screen. Radioactive signals were detected with an Amersham typhoon and quantified in AIDA software version 4.06.034.

### Immunofluorescence, click-iT EdU labeling and telomere fluorescence *in situ* hybridization (FISH)

Cells were pulse-labeled with 10 µM EdU for 30 minutes, trypsinized, washed with PBS and the cell concentration was adjusted to 0.5 million cells in 5% FBS in PBS. 100,000 or 200,000 cells were placed on the microscopic slides using cytospin centrifugation (Shandon cytospin 4; 5 minutes, 1,000 rpm) and were fixed on the slides with 4% paraformaldehyde for 15 minutes at room temperature. Cells were washed three times with PBS, permeabilized with 0.1% Triton X100, 0.02% SDS in PBS and stained with azide-coupled fluorophore (Alexa 647) after crosslinking with Click-iT chemistry. Cells were pre-blocked in 2% BSA in PBS, blocked with 10% goat serum in PBS and incubated with anti-γH2AX antibodies (1:1,000). After washing, they were incubated with secondary goat-anti mouse antibodies conjugated with Alexa 488 (1:500). After washes with PBS, cells were incubated in 4% paraformaldehyde for 5 minutes at room temperature and washed with PBS again and dehydrated with increasing concentrations of ethanol. Cy3-labeled TelC probe (PNA Bio, F1001) was diluted 1:2,000 in hybridization buffer (70% formamide, 10 mM Tris-HCl pH 7.5, 0.5% blocking reagent from Roche) applied on the slides and slides were denatured for 3 minutes at 80°C. Slides were then incubated for 3 hours in the dark at room temperature, washed twice in hybridization wash buffer 1 (70% formamide, 10 mM Tris-HCl pH 7.5) and washed 3 times in hybridization buffer 2 (0.1 M Tris-HCl 7.4, 0.15 M NaCl, 0.08% Tween-20). The second wash contained 1 µg/ml DAPI. The slides were mounted with vectashield. Images were taken with a LSM700 upright microscope at 63x magnification. γH2AX foci colocalizing with telomeric FISH signal in EdU positive and EdU negative cells were counted in FIJI and statistical analysis was done using Prism7 software.

Normal anaphases, anaphases with bridges, and micronuclei were counted in FIJI and statistical analysis was done using Prism7 software.

### Telomere fluorescence *in situ* hybridization (FISH) on metaphase spreads

Cells were treated with 0.5 µg/ml demecolcine for two hours, harvested with trypsin, incubated with 0.056 M KCl for 7 minutes at 37°C, fixed with methanol:glacial acetic acid (3:1) and stored overnight at 4°C. Cells were dropped on the microscope slides, dried overnight and fixed with 4% paraformaldehyde. Telomere FISH was conducted as described above. Images were taken with Zeiss-Axioplan and normal telomeres as well as fragile telomeres and chromatin and chromosome fusions were counted in FIJI. Statistical analysis was done in Prism7.

### RT-qPCR

RNA from 3 million cells was extracted using the Nucleospin RNA extraction kit (Macherey-Nagel). 3µg of RNA was reverse-transcribed using 200 U super Script III reverse transcriptase (Thermo Fischer), GAPDH and TERRA reverse primers for 1 hour at 55°C and the reaction was heat inactivated for 15 minutes at 70°C. 5% of the reaction was used as template for qPCR amplification with Power SYBR Green Master mix (ThermoFisher) and 0.5 µM forward and reverse primers. The program for qPCR reaction included 10 minutes at 95°C, 40 cycles at 95°C for 15 seconds and 60°C for 1 minute in an Applied Biosystems 7900HT Fast Real-Time System (Feretzaki and Lingner, 2017).

RT-qPCR for mRNAs to confirm siRNA-mediated knockdowns was performed using Luna Universal One-Step RT-qPCR Kit (NEB). 200 ng of RNA were used for one step RT-qPCR with 0.4 µM forward and reverse primers. The program for RT-qPCR included 10 minutes at 55C, 1 minute at 95°C, followed by 40 cycles at 95°C for 10 seconds and 60°C for 1 minute in an Applied Biosystems 7900HT Fast Real-Time System.

### TRAP

TRAP assay was performed as described (Kim, 1997) with modifications. 1 million cells were lysed for 30 minutes on ice in 200 µl of CHAPS lysis buffer (10 mM Tris-HCl pH 7.5, 1 mM MgCl_2_, 1 mM EGTA, 0.5% (w/v) CHAPS, 10% (w/v) glycerol, 0.1 mM PMSF, 1mM DTT) and centrifuged for 15 minutes at 4°C. Lysate concentrations were determined with the Bradford assay and adjusted to 1 µg/µl. Serial dilutions of extracts were used for the primer elongation reaction (1xTRAP buffer, 25 µM dNTPs, 2 ng TS primer, 2 µCi α-^32^P dGTP for 30 minutes at 30°C. Then telomerase was inactivated at 94°C and 2U of Taq polymerase (Eurobio) and 2 ng of ACX primers were added to the reaction. PCR included 27 cycles for 30 seconds at 94°C and 30 seconds at 60°C. Extracts treated with 0.01 M EDTA were used as negative controls. 20% of TRAP reactions were loaded on 15% non-denaturing polyacrylamide gels and separated at 200V for 4 hours. Gels were dried and exposed to a phosphorimager screen. Radioactive signals were detected with Amersham Typhoon.

### DRIP

20 million cells were harvested and each sample was split into two pellets of 10 million cells and lysed on ice with cold RLN buffer (50 mM Tris-Cl, pH 8.0; 140 mM NaCl; 1.5 mM MgCl_2_; 0.5% v/v Nonidet P-40; 1 mM DTT and 100 U/ml RNasin Plus). After centrifugation (300 *g*, 2 minutes, 4°C), the nuclei-enriched pellets were resuspended in 500 µL of RLT buffer with 0.13% β-mercaptoethanol and homogenized by passing through a 0.9×40mm needle. Nucleic acids were then isolated by a phenol:chloroform:isoamylalcohol (25:24:1; Biosolve BV) extraction, followed by isopropanol precipitation. The two nucleic acids pellets coming from the same sample were joined together and dissolved in 300 µl H_2_O. Nucleic acid solutions were sonicated using a Focused-ultrasonicator (Covaris E220) to obtain fragments below 300 bp. 150 µl of sonicated nucleic acids were treated for 90 minutes at 37°C with either 10 µl of RNase H (1 U/µl; Roche) or H_2_O. The reaction was stopped by addition of 2 µl of 0.5 M EDTA. 50% protein G sepharose beads slurry (GE Healthcare Life Sciences) was blocked with 1mg/ml yeast tRNA for 1 hour at 4°C. The samples were then diluted with 1,200 µl of buffer 1 (10 mM Hepes-KOH, pH 7.5; 275 mM NaCl; 0.1% SDS; 1% Triton X-100; and 0.1% Na-deoxycholate) and precleared for 1 hour at 4°C. 650 µl of precleared extract were used per immunoprecipitation reaction together with 80 µl of blocked 50% protein G beads slurry, and 1 μg of S9.6 antibody (022715; Kerafast) or mouse IgG (sc-2025; Santa Cruz Biotechnology). The reactions were incubated for 2 hours at 4°C on a rotating wheel and then washed consecutively for 5 minutes each time with 1 ml of buffer 2 (50 mM Hepes-KOH pH 7.5; 140 mM NaCl; 1% Triton X-100; 0.1% Na-deoxycholate; and 1 mM EDTA), buffer 3 (50 mM Hepes-KOH pH 7.5; 500 mM NaCl; 1% Triton X-100; 0.1% Na-deoxycholate; and 1 mM EDTA), buffer 4 (10 mM Tris–HCl pH 8.0; 250 mM LiCl; 1% NP-40; 1% Na-deoxycholate; and 1 mM EDTA), and 1× TE buffer (10 mM Tris–HCl pH 8.0, and 1 mM EDTA). Beads and inputs were incubated with reverse crosslinking buffer (1%SDS, 0.1M NaHCO_3_, 0.5mM EDTA pH 8.0, 20 mM Tris-HCl pH 8.0, RNase-DNase free, Roche) for 1 hour at 65°C and DNA was extracted using QIAquick PCR purification kit. Samples were transferred to the Hybond N+ membrane using dot bot and UV-crosslinked before hybridization with telomeric probe as described above.

### Cell cycle

Cells were incubated with 10 µM EdU for 30 minutes, harvested and washed with PBS. Cells were fixed with 70% ethanol at a concentration of 1 million cells per mL. The next day cells were washed with PBS, permeabilized with 0.5% Triton X-100 for 20 minutes, stained with Alexa-488 conjugated azide using Click-iT chemistry for 30 minutes in the dark and with 1µg/mL DAPI, transferred to FACS tubes and cell cycle profiles were acquired with Gallios (Beckman Coulter). Further analysis was done with Kaluza software.

### C-circle assay

The C-circle assay was performed as described before (Henson et al., 2009). Briefly, 300 ng of genomic DNA was digested with 4 U/μg (each) HinfI and RsaI (NEB) in buffer 2 and 10 μg/ml RNase A for 1 hour at 37°C. 30 ng, 10 ng, 3 ng and 1 ng of digested DNA was taken to perform the φ29 reaction which contained in 20 μl 0.2 mg/ml BSA, 0.1% Tween, 1 mM each of dATP, dGTP, dTTP, 1xφ29 buffer and 7.5 U φ29 DNA polymerase). The mixture was incubated for 8 hours at 30°C and ϕ29 DNA polymerase was inactivated by incubation at 65°C for 20 minutes. The reaction products were dot-blotted on a positively charged nylon membrane and UV-crosslinked. The membrane was prehybridized at 50°C for 1 hour in Church buffer and hybridized overnight at 50°C with a ^32^P-radiolabeled telomeric probe. The membrane was washed three times with 4xSSC at RT and once with 4xSSC and 0.1% SDS at 50°C (30 minutes each wash) and exposed to phosphorimager screen. Radioactive signals were detected with Amersham typhoon and quantified in AIDA software version 4.06.034.

### 2-step QTIP

Telomeric chromatin precipitation was performed as described before (Majerska, 2017) with a number of modifications. SILAC labeling was not used. 1 billion cells were used per IP reaction. After harvesting, cells were washed twice with PBS, fixed in 1% formaldehyde and 2 mM EGS for 15 minutes at 25°C at a cell concentration of 10 million cells/ml. For chromatin enrichment cells were resuspended in the chromatin enrichment buffer (50 mM Tris-HCl pH 8.0, 1% SDS, 1 mM EDTA) at 10 million cells/ml, incubated on the wheel for 5 minutes at 25°C and centrifuged at 4,000 rpm for 5 minutes. Chromatin enriched pellets were resuspended at 20 million cells/ml in 4 batches of 25 ml of LB3 buffer (10 mM Tris–HCl, pH 8.0; 200 mM NaCl; 1 mM EDTA–NaOH, pH 8.0; 0.5 mM EGTA–NaOH, pH 8.0; 0.1% w/v sodium deoxycholate; 0.25% w/v sodium lauryl sarcosinate; and EDTA-free protease inhibitor complex (Roche)) and sonicated using a Branson Tip sonicator (30% power, 10 seconds constant pulse, and 20 seconds pause for a total sonication time of 9 minutes). Sonicated extracts were dialyzed against immunoprecipitation (IP) buffer (50 mM Tris-HCl pH 8.0, 600 mM NaCl, 10 mM EDTA pH 8, 0.75% Triton X-100) and a first IP was performed using either ANTI-FLAG M2 Affinity Agarose Gel (Sigma) or mouse IgG coupled to sepharose beads that had been blocked with yeast tRNA (1mg/mL) for 1 hour at 4°C. 1.6 ml of 50% beads slurry was used per IP, reactions were performed overnight at 4°C. After 5 rounds of washes with IP buffer for 5 minutes each, five rounds of elutions were performed with 100 µg/mL of FLAG peptide (Sigma). A second IP was performed overnight at 4°C with home-made affinity-purified anti-TRF1 and anti-TRF2 antibodies coupled to protein G sepharose beads that had been blocked with yeast tRNA. 0.8 ml of 50% beads slurry was used per IP. After washes with buffer 1 (0.1% SDS, 1% Triton X-100, 2 mM EDTA pH 8.0, 20 mM Tris-HCl pH 8.0, 300 mM NaCl), buffer 2 (0.1% SDS, 1% Triton, 2 mM EDTA pH 8.0, 20 mM Tris-HCl pH 8.0, 500 mM NaCl), buffer 3 (250 mM LiCl, 1% NP-40, 1% Na-Deoxycholate, 1 mM EDTA pH 8.0, 10 mM Tris-HCl pH 8.0) and buffer 4 (1 mM EDTA pH 8.0, 10 mM Tris-HCl pH 8.0) beads were dried and resuspended in 2.5 bed bead volume equivalent of 0.25 M ammonium hydroxide for elution for 15 minutes at 37°C. Four rounds of elution were performed, eluates were pooled, flash-frozen and lyophilized.

### Protein Digestion

Lyophilized samples were digested following a modified version of the iST method (Kulak et al. 2014). Briefly, pellets were resuspended in 100 µl miST lysis buffer (1% sodium deoxycholate, 100 mM Tris-HCl pH 8.6, 10 mM DTT) by vigorous vortexing. An internal standard protein (alcohol dehydrogenase from *S.cerevisiae*, Merck-Sigma A7011, 0.02 µg) was spiked in all samples, while 0.05 µg of bovine beta-lactoglobulin (Merck-Sigma L3908) were added only to the TRF1/2 QTIP immunoprecipitates in order to quantify TMT ratio compression effects. Resuspended samples were heated at 95°C for 30 minutes to denature proteins and reverse formaldehyde-induced crosslinking. Samples were then diluted 1:1 (v:v) with water containing 4 mM MgCl_2_ and Benzonase (Merck-Novagen 70746-4, 1:100 of reaction volume = 51.6 Units) and incubated for 1 hour at RT to digest nucleic acids crosslinked to proteins. Reduced disulfides were alkylated by adding ¼ vol (50 µl) of 160 mM chloroacetamide (final 32 mM) and incubating at RT for 45 minutes in the dark. Samples were adjusted to 3 mM EDTA and digested with 1 µg Trypsin/LysC mix (Promega V5073) for 1 hour at 37°C, followed by a second overnight digestion with a second, identical aliquot of proteases. To remove sodium deoxycholate, digests were phase extracted by adding 600 µl of ethyl acetate containing containing 1% TFA, vortexing for 2 minutes and centrifugation. The bottom aqueous phase containing peptides was collected, diluted 2x with 0.5% formic acid and desalted on strong cation exchange (SCX) cartridges (SOLA Cartridges (Thermo Fisher: 60209-002)) by centrifugation. After washing the SCX cartridges with 0.5% formic acid, 20% acetonitrile, peptides were eluted in 200 µl of 80% MeCN, 19% water, 1% (v/v) ammonia).

An aliquot of 1/15 of samples from experiment replicate 1 (2-2) was analyzed directly by LC-MS and label-free quantitation to assess the quality of the digestion, the specificity of the preparation and, by comparison with known amount of a standard HeLa digest, obtain an approximate estimate of the total amount (by mass) of peptides present before TMT labelling. The samples appeared to contain approximately 30 µg of material, with antibody chains used for purification seemingly quantitatively dominant, thus making up the bulk of protein material.

### TMT labelling

Eluates after SCX desalting were dried, resuspended in 100 µl water and dried again to remove excess ammonia. Digests were then resuspended in 40 µl of 50 mM TEAB buffer, pH 8.0 and reacted with 0.4 mg of TMT reagent for 1hour at RT, after which excess reagent was quenched with 8ul of 50% hydroxylamine for 20 minutes at RT. For each replicate experiment, untreated (NT, day 0) and KD samples (day 4, day 7) were mixed with their corresponding negative IgG controls in a six-plex TMT experiment using, respectively, TMT channels 126, 127N, 128N, 129N, 130N and 131. Neutron-encoded C labels were not used). TMT multiplex samples were dried and desalted on C18 SepPak well plates. An aliquot (5%) was injected before fractionation to assess labelling completion (>98% peptide spectrum matches) by database search with TMT as variable modification (MASCOT software).

### Peptide fractionation

Dried desalted eluates were dissolved in 4M Urea containing 0.1% Ampholytes pH 3-10 (GE Healthcare) and fractionated by off-gel focusing as described (Geiser et al., 2011). The 24 peptide fractions obtained were desalted on a SepPak microC18 96-well plate (Waters Corp., Milford, MA), dried and redissolved in 30 µl of 0.05% trifluroacetic acid, 2% (v/v) acetonitrile for LC-MS/MS analysis.

### MS analysis

Data-dependent LC-MS/MS analysis of extracted peptide mixtures after digestion was carried out on a Fusion Tribrid Orbitrap mass spectrometer (Thermo Fisher Scientific) interfaced through a nano-electrospray ion source to a RSLC 3000 HPLC system (Dionex). Peptides were separated on a reversed-phase custom packed 40 cm C18 column (75 μm ID, 100Å, Reprosil Pur 1.9 um particles, Dr. Maisch, Germany) with a 4-76% acetonitrile gradient in 0.1% formic acid (total time 140 minutes). Full MS survey scans were performed at 120’000 resolution. In data-dependent acquisition controlled by Xcalibur 4.0.27 software (Thermo Fisher Scientific), a data-dependent acquisition method was used that optimized the number of precursors selected (“topspeed”) of charge 2+to 5+ while maintaining a fixed scan cycle of 1.5 second. The ion isolation window used was 0.7 Th.

Peptides were fragmented by higher energy collision dissociation (HCD) with a normalized energy of 35%. MS2 scans were done at a resolution of 30’000 in the Orbitrap cell, which is sufficient to resolve 6-plex TMT reporter ions. The *m/z* of fragmented precursors was then dynamically excluded from selection during 60 seconds.

### MS data analysis

Data files were analysed with MaxQuant 1.6.3.4 (Cox and Mann, 2008; Cox et al., 2011) incorporating the Andromeda search engine (Cox et al., 2011). Cysteine carbamidomethylation and TMT labelling (peptide N-termini and Lysine side chains) were selected as fixed modification while methionine oxidation and protein N-terminal acetylation were specified as variable modifications. The sequence database used for searching was the human Reference Proteome based on the UNIPROT database (www.uniprot.org), version of October 29^th^, 2017, (2017_10) containing 71’803 sequences. This was completed with custom databases containing most usual environmental contaminants (keratins, trypsin, etc), the sequence of yeast ADH and a collection of mouse (554) and sheep (37) sequences of immunoglobulins (also from UNIPROT). Mass tolerance was 4.5 ppm on precursors (after recalibration) and 20 ppm on HCD fragments. Both peptide and protein identifications were filtered at 1% FDR relative to hits against a decoy database built by reversing protein sequences. For unlabelled samples, iBAQ values (Schwanhäusser et al., 2011) generated by MaxQuant in label-free quantitation were used. For TMT analysis, the raw reporter ion intensities generated by MaxQuant and summed for each protein group were used in all following steps to derive quantitation. Only identified peptide ions with a precursor intensity fraction (PIF parameter) greater than 0.75 were accepted and used for TMT quantitation.

### Processing of quantitative data and statistical tests

The MaxQuant output table “proteinGroups.txt” for the three TMT replicate samples was first processed to remove proteins matched to the contaminants database as well as proteins identified only by modified peptides and reverse database hits, yielding a first unfiltered list of 2674 identified proteins. As the composition of positive and negative samples was very different, no global normalization was applied to the data. After log-2 transformation of all intensity values, a plot of the intensity of the spiked-in control (yeast ADH) showed a slight decrease of signal intensity correlating with the order of MS injection (Figure S4). A global correction factor was applied per TMT sample mix to compensate for this effect, which is probably in partly due to gradual deterioration of the performance of the LC column over the >80h of continuous MS measurement. The larges correction factor was +0.28 (log2 scale) relative to the median of all samples. Next, the table was filtered to retain only proteins identified by a minimum of two “razor or unique” peptides and having at least 12 out of 18 total quantitative values (1695 proteins left). Ratios (difference in log scale) were calculated for each sample relative to its IgG control.

A one-sample T-test (p-value filter at 0.05) was performed on all values by comparing the 9 TERF1/2 IP’s vs 9 IgG controls to identify a set of invariant proteins that did not significantly change in intensity between IgG controls and positive IP’s. 249 proteins (including many keratins and common contaminants) passed the test and were removed from the dataset, leaving 1446 protein groups. The obtained ratio QTIP/IgG was recorded in the tables as “ specificity ratio”. To determine proteins significantly changing between NT and the knockdown (day4, day7) conditions, we applied a paired T-test (each sample against its IgG control) with Benjamini-Hochberg FDR correction (Benjamini and Hochberg, 1995) and threshold at 0.05 on the adjusted p-value. No protein passed the test at day 4 while 157 proteins passed the test at day 7.

### Raw data deposition

All raw MS data toghether with MaxQuant output tables are available via the Proteomexchange data repository () with the accession number PXD016826 (Username; Password: ifxrqkd2).

---

**Figure S1 (related to Figure 1).**
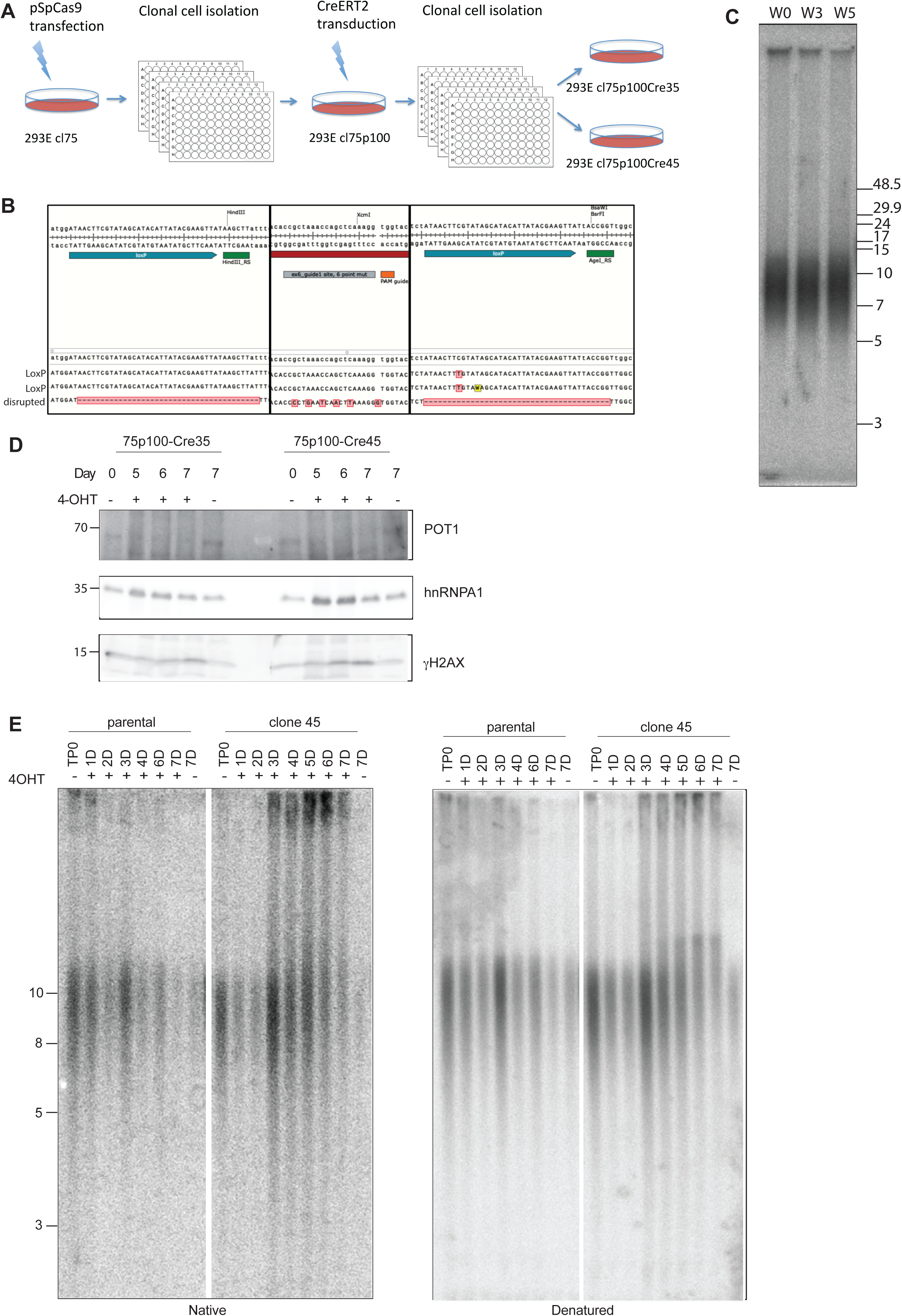
(A) Schematic for generation of POT1 KO cell lines. (B) Genotype of the clone 75p100. Two alleles have successfully integrated loxP sites, the third allele has one base insertion leading to the frame shift and lack of the WT protein. (C) Telomere length analysis for clone 75p100. (D) Western blot for clones 35 and 45 upon POT1 removal. (E) Timecourse for telomere length analysis for parental (clone 75) and LoxP containing clone 45 cell lines upon Cre induction with 0.5 µM of 4-OHT.

**Figure S2 (related to Figure 1).**
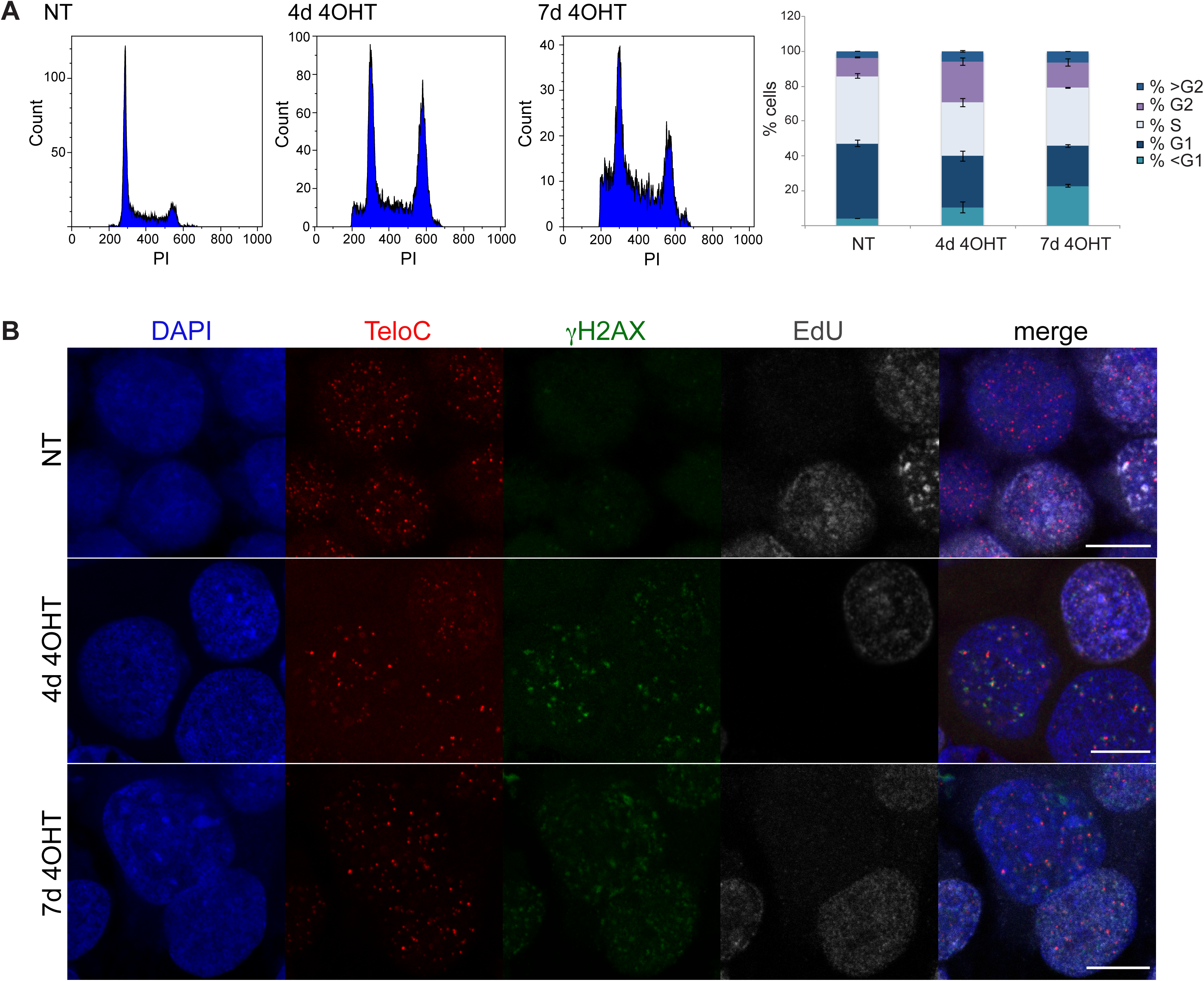
(A) Cell cycle profiles and quantification for non-treated (NT) clone 35 cells and clone 35 treated with 4-OHT for 4 and 7 days. Graph represents data from three independent experiments. (B) Representative pictures of TIF assay in EdU labeled cells. Scale bar equals 5 µm.

**Figure S3 (related to Figure 3).**
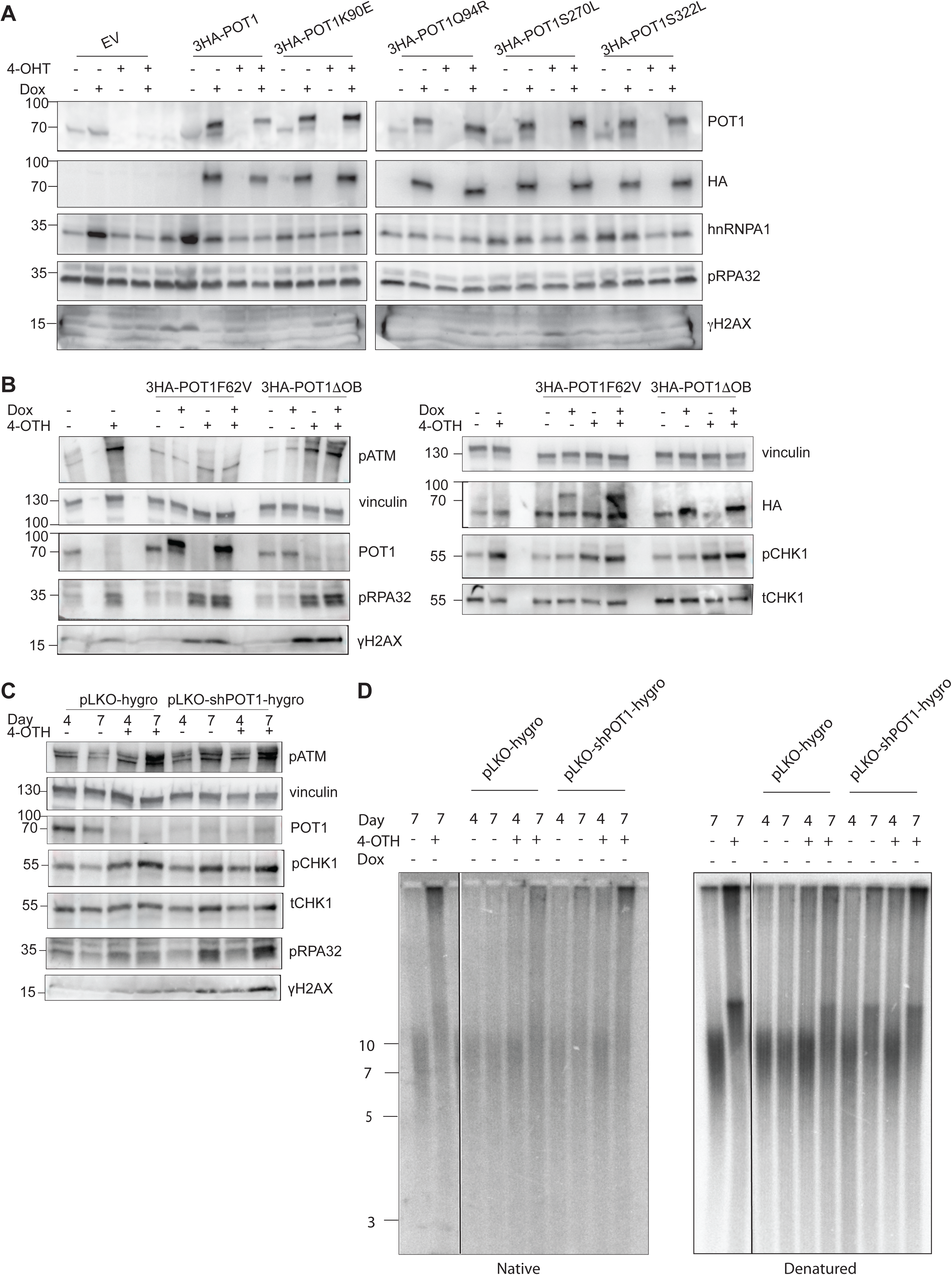
(A), (B) Western blot for POT1 and DDR markers upon POT1 removal in clone 35 and overexpression of ectopic POT1 mutants. Cells were induced for 7 days with 0.5 µM 4-OHT and for 4 days with 1µg/mL doxycycline. (C) Western blot for POT1 and DDR markers upon POT1 removal in clone 35 using either Cre recombinase or knockdown with lentivirus containing shRNA against POT1. (D) Telomere length and G-overhang length analysis for clone 35 upon POT1 removal in clone 35 using either Cre recombinase or knockdown with lentivirus containing shRNA against POT1.

**Figure S4 (related to Figure 4).**
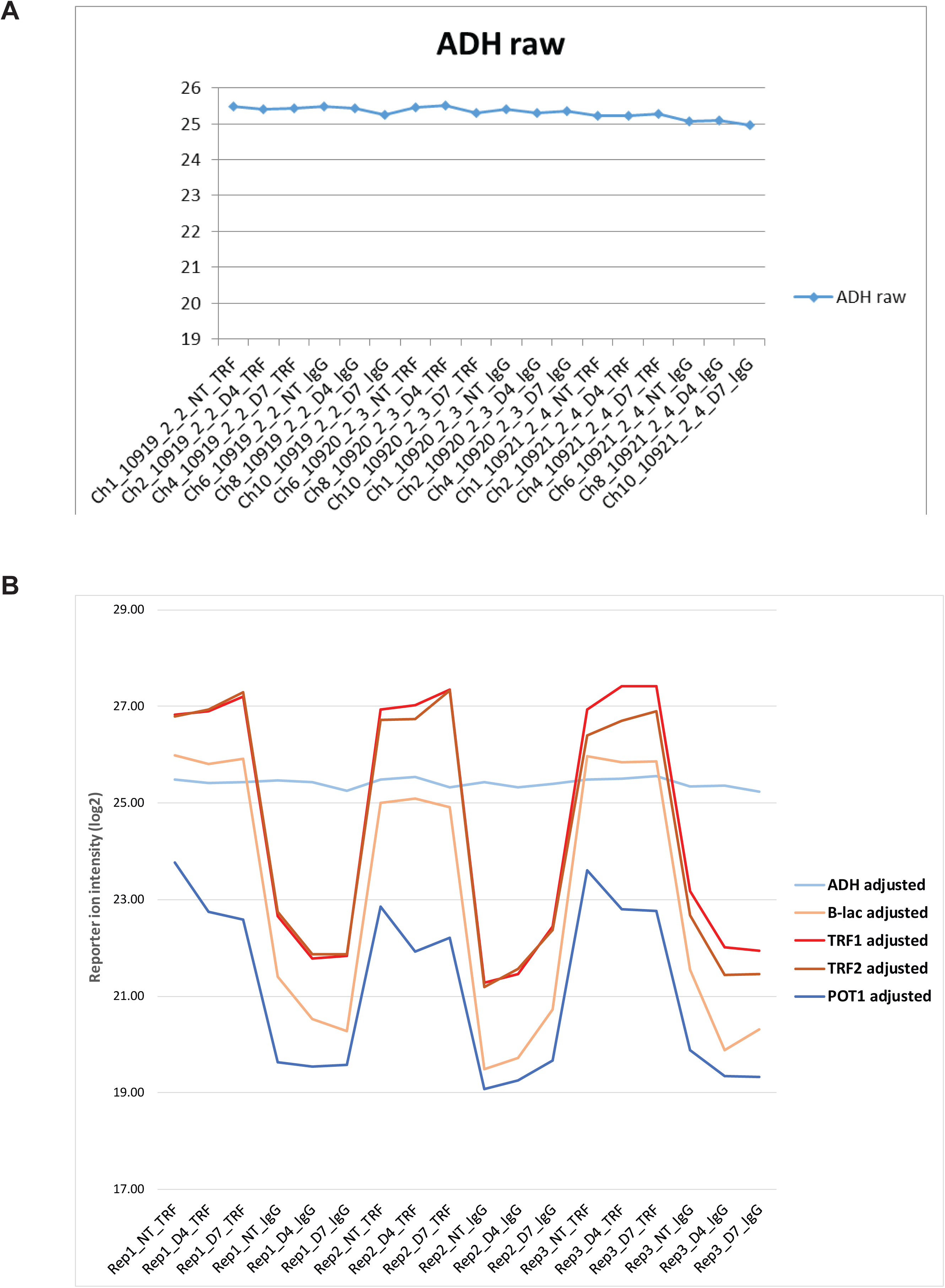
(A) Yeast Alcohol dehydrogenase (ADH, UNIPROT P00330) standard was spiked in all samples in equal amounts. MS analysis was done in the order 2-2, 2-3, 2-4 shown here. (B) TERF1/2 and standards intensities after adjustment by the spiked-in standard (yeast ADH). Bovine beta-lactoglobulin (B-lac) was spiked uniquely in Q-TIP samples to quantify the maximal ratio measurable by TMT in the conditions of the experiment. B-lac as well as TERF1/2 had average (QTIP/IgG) differences of approximately 5.0 (log2), corresponding to a fold change of 32x in a linear scale. A preliminary label-free MS analysis of the unfractionated samples of replicate 1 showed that shelterins were essentially undetectable in the IgG controls, confirming this evaluation (Supplementary Table 1). Measurable TMT ratios are also impacted by the absolute signal intensity, whereby less abundant proteins are expected to have lower maximal measurable ratios.

**Figure S5 (related to Figure 6).**
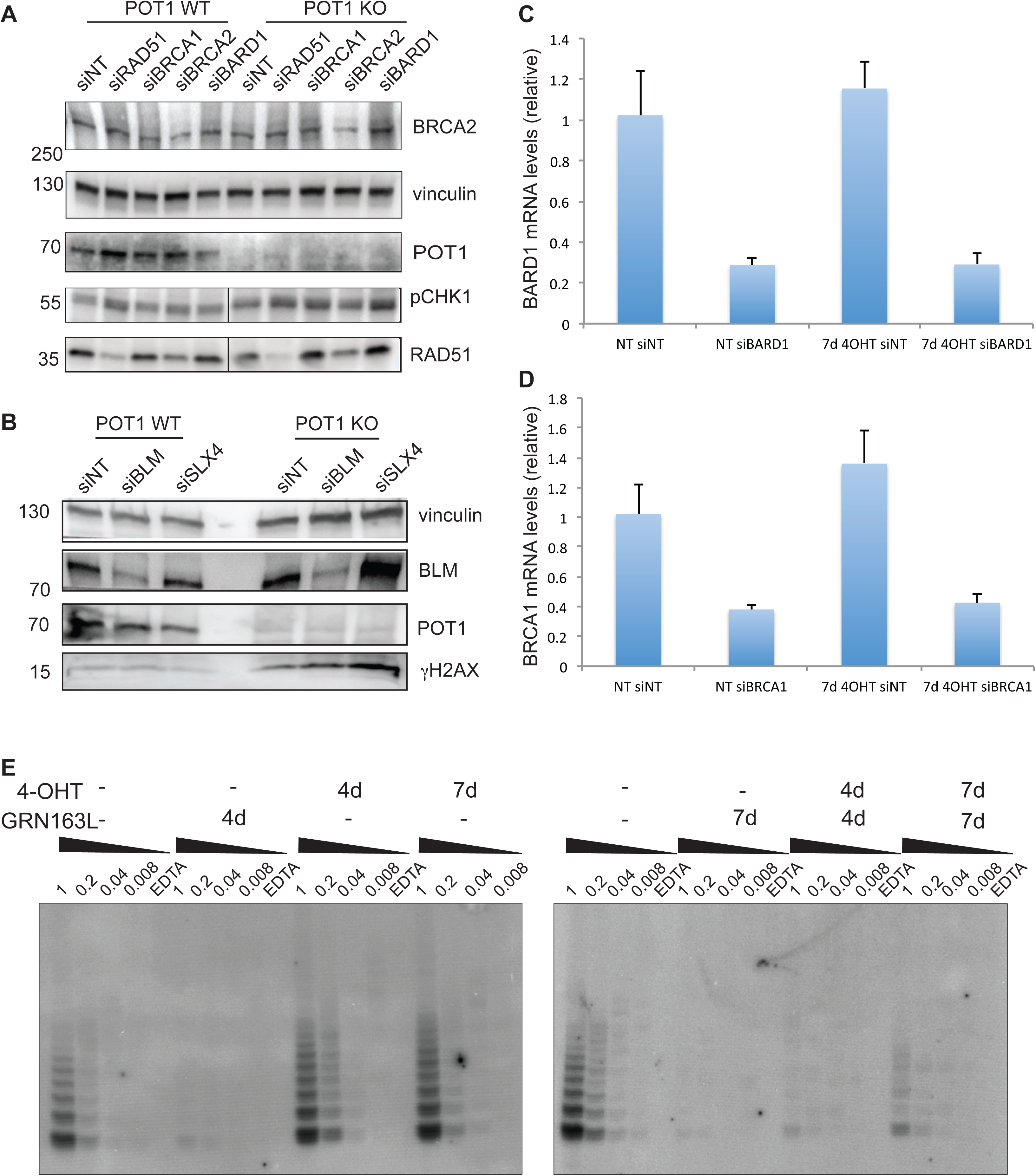
(A) Telomere repeat amplification protocol (TRAP) assay for clone 35 treated with 0.5 µM 4OHT and/or 1µM GRN163L. As negative controls, EDTA-containing extracts were run in parallel. (B) Western blot for POT1, DDR markers and recombination proteins in clone 35 upon depletion of selected recombination proteins with siRNA and removal of POT1. (C) qPCR to confirm depletion of BRCA1 and BARD1 in clone 35 upon depletion of these proteins with siRNA. (D) Western blot for POT1, γH2AX BLM in clone 35 upon depletion of BLM with siRNA and removal of POT1. All experiments were performed in triplicate.

**Figure S6 (related to Figure 6).**
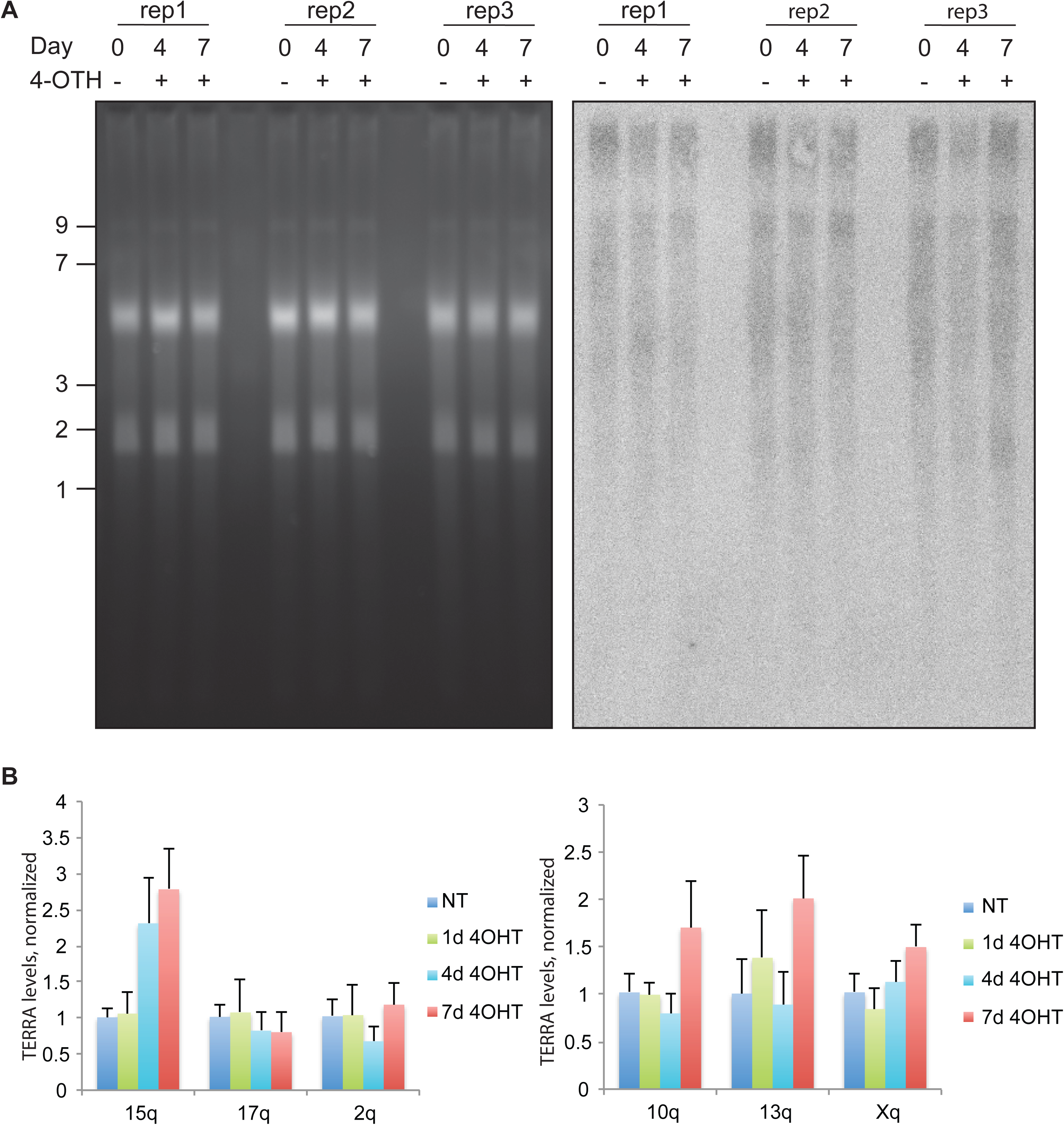
(A) qPCR to detect selected TERRAs in clone 35 upon POT1 removal. (B) Northern blot to detect total TERRA levels in clone 35 upon POT1 removal.

